# DNA glycosylases Ogg1 and Mutyh mediate gene expression of PRC2 target genes important for neuronal processes underlying memory formation

**DOI:** 10.1101/2024.04.16.589756

**Authors:** Andreas Abentung, Teri Sakshaug, Rabina Dumaru, Nina-Beate Liabakk, Mingyi Yang, Junbai Wang, Magnar Bjørås, Katja Scheffler

## Abstract

Base excision repair (BER) initiated by DNA glycosylases is known to preserve genomic integrity by removing damaged bases. Recently, several DNA glycosylases were identified as potential readers of epigenetic modifications and proteins involved in BER have been associated with active DNA demethylation. DNA glycosylases Ogg1 and Mutyh were shown to alter the hippocampal transcriptome associated with cognitive function and independent of global DNA damage accumulation. However, the mechanism of DNA glycosylases in regulating cognition and their role in epigenetic remodeling in the brain remains elusive. Here we report that the combined deficiency of Ogg1 and Mutyh impairs spatial but not associative long-term memory. We demonstrate that Ogg1 or Mutyh modulate DNA methylation at gene regulatory regions of polycomb repressive complex 2 (PRC2) target genes in the adult hippocampus. Moreover, we find that the distribution of the PRC2 complex and histone modifications associated with PRC2 activity changes in both hippocampal neurons and glia depend on Ogg1 and Mutyh. Epigenetic alterations correlated with cell-type specific gene expression changes which were associated with pathways important for neuronal function and cognition. Our results provide a novel role for Ogg1 and Mutyh beyond DNA repair in modulating the epigenome to control transcriptional responses in the brain important for memory formation.

## Introduction

The canonical functions of DNA repair play an essential role in maintaining genomic integrity and stability across organisms^1^. Genomic instability caused by an accumulation of DNA damage is a hallmark of aging^2^ and has been linked to DNA mutations, altered gene expression and cognitive impairment in the human brain^3,4^. Due to its high metabolic rate, high transcriptional activity and long lifespan, brain cells are highly susceptible to DNA damage^5^. Consistently, mutations in DNA repair genes and impaired DNA repair pathways are associated with pre-mature aging and various neurodegenerative diseases^6,7^. One of the major threats challenging genome integrity are reactive oxygen species (ROS)^8^. ROS can arise as a byproduct during normal cellular metabolism and can inflict oxidative DNA damage^9^. Given its low redox potential, the DNA base guanine is particularly susceptible to oxidation, leading to the formation of the highly pre-mutagenic base lesion 8-oxoguanine (8-oxoG)^10^. 8-oxoG stands out as one of the most well studied oxidative DNA lesions, capable of inducing point mutations owing to its tendency to miscode. This miscoding potential prompts DNA polymerases to favor the insertion of Adenine (A) opposite 8-oxoG instead of the correct Cytosine (C). If left uncorrected, these A:8-oxo-G mispairs can lead to C:G → A:T transversion mutations^11^. 8-oxoG DNA glycosylase 1 (Ogg1) recognizes and facilitates the removal of 8-oxoG within the genome through the base excision repair pathway^12,13^. Adenine DNA glycosylase (Mutyh) excises adenine mis-incorporated opposite of 8-oxoG during replication allowing Ogg1 to remove 8-oxoG after insertion of the correct base^14^.

Emerging evidence indicates that DNA glycosylases may have additional functions beyond canonical DNA repair by altering the epigenetic landscape and thereby regulating transcription^15^. In the mouse brain, Ogg1 and Mutyh regulate neuronal gene expression related to learning and memory independent of global accumulation of 8-oxoG^16^. Consistent with these findings, catalytic inactive Ogg1 facilitates pro-inflammatory gene expression emphasizing an important role of DNA glycosylases independent of 8-oxoG repair^17^. Interestingly, several DNA glycosylases were identified as potential readers of epigenetic DNA modifications^18,19^ and have been implicated in active DNA demethylation^20,21^. Additionally, DNA glycosylases have been shown to interact with several epigenetic modifiers such as DNA methyltransferases^22,23^, histone demethylases^24^ and members of the polycomb repressor complex (PRC)^25^.

It has been shown that 8-oxoG, located in promoter regions, itself has the potential to elicit epigenetic-like properties by modulating gene transcription^26,27^. However, recent evidence suggests that this occurs mainly independent of its repair and that DNA glycosylases Ogg1 and Mutyh modulate gene transcription without altering the genomic distribution of 8-oxoG^28^. However, the molecular mechanisms through which DNA glycosylases impact gene transcription in the brain contributing to learning and memory still remain elusive.

In this study, we investigate the impact of Ogg1 and Mutyh on the epigenetic landscape in the adult mouse hippocampus. We show that loss of DNA glycosylases affects spatial but not associative long-term memory. Mechanistically, we find that Ogg1 and Mutyh modulate DNA methylation level at regulatory regions of PRC2 target genes and histone modifications associated with PRC2 activity in both neurons and glia. Taken together, our results indicate a novel role for Ogg1 and Mutyh beyond canonical DNA repair in modulating the epigenome to control transcriptional responses important for memory formation in different brain cells.

## Materials and Methods

### Animals

All experiments were approved by the Norwegian Animal Research Authority (NARA) and conducted in accordance with the laws and regulations controlling experimental procedures in live animals in Norway and the European Union’s Directive 86/609/EEC. Mice were housed and bred at the Comparative Medicine Core Facility (Trondheim, Norway) in a 12-hour light/dark cycle with access to food and water *ad libitum*. Housing was under standard laboratory conditions of 19-22°C with air humidity of 50-60%. Ogg1 and Mutyh deficient C57BL/6N mice were used for all the experiments included in this study. Double knock out (DKO) mice were obtained by crossing *Ogg1^-\-^* and *Mutyh^-\-^* mice. Wild type (WT) C57BL/6N male mice were used as control groups. For all experiments 6 months old male mice were used. Mouse litters were genotyped using genomic DNA obtained from ear biopsies. All knock out strains have been previously described elsewhere^16^.

### Contextual fear conditioning

Mice were handled 5 minutes for three days prior to the experiment. Contextual fear conditioning was performed in a 17 x 17 x 25 cm chamber with transparent walls and a metal rod floor, cleaned with soap-water and illuminated to 150 lux (Ugo Basile, Fear Conditioning Animal System, 46001) as previously described^29^. After a 120s acclimation period, mice were conditioned with three presentations of a 0.60 mA scrambled foot shock, with a 120s inter-shock interval. The mice were allowed to remain in the chamber for an additional 120s following the last stimulus presentation. Short-term and long-term fear memories were assessed by re-exposing the mice for 300s either 1 hour or 24 hours later to the conditioning chamber without the administration of a foot shock. Freezing was measured as an index of fear^30^ defined as no visible movement except that required for respiration. Freezing time was scored and analyzed with AnyMaze software and converted to a percentage [(duration of freezing /total time) × 100].

### Flinch-jump test

Reactivity to the foot shock was evaluated in the same apparatus used for contextual fear conditioning. After a 120s acclimation period, mice were subjected to a series of 1 second foot shocks with gradually increasing amperage (0.1 mA every 30 seconds) starting from 0.1 mA. Mice were scored for their first visible response to the shock (flinch), their first pronounced motor response (run or jump) and their first vocalized distress, as previously described^31^.

### Novel object location

The animals were quartered in a 12-hour reversed light/dark cycle. Prior to training, mice were handled 5 minutes for five consecutive days and then habituated to the experimental apparatus (50 x 50 x 30 cm open field area) for 5 minutes for three consecutive days in the absence of objects as previously described^29^. During training, mice were placed into the experimental apparatus containing two identical objects (Lego blocks) and allowed to explore for 10 minutes. Short-term and long-term retention tests were conducted by placing the mice back into the experimental apparatus for 10 minutes either 1 hour or 24 hours after training. To assess spatial object location-dependent memory, the placement of one familiar object remained unchanged whereas the other copy of the familiar objects was moved and placed adjacent to a different corner inside the experimental apparatus. Exploration was scored when the mouse’s head entered the area adjacent to the object. All training and testing trials were videotaped and analyzed with AnyMaze software. The relative exploration time (t) is expressed as a percent discrimination index (D.I. = tnovel / (tnovel + tfamiliar) × 100%). Mean exploration times were then calculated and the discrimination indices between experimental groups were compared. Animals that explored each object for less than 3 seconds during training or explored both objects for less than 10 seconds total during testing were excluded from the analysis.

### Tissue collection

Animals were sacrificed by cervical dislocation and hippocampi were dissected, isolated, snap-frozen in liquid nitrogen and stored at −80°C until further use. All mice included in high throughput sequencing experiments were re-genotyped by using genomic DNA obtained from tail biopsies after being sacrificed.

### DNA isolation

Total DNA was extracted from snap-frozen hippocampi by using AllPrep DNA/RNA/Protein mini Kit (Qiagen, 80004), according to manufacturer’s protocol. Frozen tissue was transferred to tubes with 1.4 mm stainless steel beads containing 350 ml buffer RLT and homogenized using a MagNA Lyser Rotor (5000 rpm, 10sec, Roche). Samples were further processed according to the respective protocols. DNA concentrations were measured by using a ND-1000 spectrophotometer (Nanodrop Technologies).

### Whole genome bisulfite-sequencing

Total DNA from two replicates each genotype was sent to BGI Tech Solutions Co., Hong Kong. Bisulfite conversion, library preparation and DNA sequencing was performed by BGI as previously described^32^. Briefly, total hippocampal DNA was fragmented by sonication using a Bioruptor (Diagenode) to a mean size of approximately 250 bp, followed by blunt-ending and dA addition to 3’-end. Finally, adaptor ligation (in this case of methylated adaptors to protect from bisulfite conversion) was performed according to the manufacturer’s instructions. Ligated DNA was bisulfite converted using the EZ DNA Methylation-Gold kit (ZYMO). After treatment with sodium bisulfite, unmethylated cytosine residues are converted to uracil whereas 5-methylcytosine (5mC) remains unaffected. Different insert size fragments were excised from the same lane of a 2% TAE agarose gel.

Products were purified by using QIAquick Gel Extraction kit (Qiagen) and amplified by PCR. After PCR amplification, uracil residues are converted to thymine. Pooled libraries were sequenced (paired-end, 100 bp in length) on a BGISEG-500 sequencer with an average depth of 1.3 billion reads per sample.

### Sorting of cell type-specific nuclei

Hippocampal nuclei were extracted, isolated and sorted with some minor modifications as previously described^33^. In brief, snap-frozen hippocampi were pooled from 5 animals of the same genotype in 1 ml of low sucrose buffer (0.32 M sucrose, 10 mM HEPES pH 8.0, 5 mM CaCl2, 3 mM Mg(CH3COO)2, 0.1 mM EDTA, 0.1% Triton X-100, 1 mM DTT, 1x Complete EDTA free protease inhibitor cocktail (Roche, 11873580001) and 20U/ml of RNase Inhibitor (Applied Biosystems, N8080119)). All steps except the crosslinking were performed at 4°C or on ice. Hippocampal tissue was homogenized by applying pestle “loose” and “tight” 40 times, respectively. For nuclei fixation, formaldehyde was added at a final concentration of 1% and the samples were incubated at room temperature on a rotator for 10 minutes. The crosslinking reaction was stopped by adding glycine to a final concentration of 125 mM for 5 minutes at room temperature. Nuclei were then pelleted by centrifugation, re-suspended and washed twice with 1 ml of low sucrose buffer. Subsequently, nuclei were purified through a sucrose cushion (10 mM HEPES pH 8, 1 M sucrose, 3 mM Mg(CH3COO)2, 1 mM DTT, 6 ml of cushion for 1,5 ml of lysate) by centrifugation at 3200 g for 10 minutes in a 15ml Falcon tube. Afterwards, nuclei were re-suspended in nuclei buffer (PBS, 0,1% Tween 20 in PBS, 5% BSA, 1x Complete EDTA free protease inhibitor cocktail (Roche, 11873580001) and 20U/ml of RNase Inhibitor (Applied Biosystems, N8080119)) containing 3% goat serum and cleared by filtering through a 40 μm cell strainer. The nuclei suspension was stored on 4°C overnight. On the next day, nuclei were stained with anti-NeuN mouse antibody (Millipore, mab377) diluted 1:500 for 60 minutes at 4°C. The samples were washed twice with nuclei buffer containing 3% goat serum. Subsequently, the samples were stained for 60 minutes at 4°C with secondary antibody (Alexa 488, A11001, Life Technologies) diluted 1:1000. Nuclei were then washed twice with nuclei buffer and stored until sorting. Directly before the sorting, nuclei were dissociated by passing them 10 times through a 25G needle and by filtering through a 40 μm cell strainer into ice-cold conical tubes containing 1.8 ml of filtered PBS-BSA (5%). Lastly, 2 μl of DyeCycle Ruby was added to the nuclei suspension. Nuclei populations were selected using a FACSAria II (BD Bioscience) for DyeCycle Ruby staining, singlet nuclei and NeuN+ positive staining. Both NeuN stained (NeuN+) and unstained (NeuN−) fractions were collected. The average purity of the sorted nuclei exceeded 95%, yielding highly cell type-specific material. The NeuN positive population contains every neuron expressing NeuN endogenously, which are primary excitatory neurons as well as interneurons. NeuN negative cells consist to a very large percentage of glial cells but also contain other cell types. Flow cytometry data were analyzed and plotted by using FlowJo (BD Bioscience).

### Chromatin Immunoprecipitation (ChIP)

Sorted nuclei were pelleted by centrifugation at 3,200 g for 15 minutes, transferred to Diagenode shearing tubes and carefully re-suspended in RIPA buffer (10 mM Tris-Cl, pH 8.0, 140 mM NaCl, 1 mM EDTA, 1% Triton X-100, 0.1% sodium deoxycholate, 1% SDS and 1x Complete EDTA free protease inhibitor cocktail (Roche, 11873580001)). Samples were incubated 10 minutes at 4°C and then sheared using a Bioruptor (Diagenode, Belgium) for 4 times at 5 cycles and followed by 2 cycles at 30″ ON/OFF high power. Samples were spun down in between every 5 cycles. Sheared chromatin was cleared by centrifugation at 10,000 rpm for 10 minutes and the supernatant was stored in DNA low-binding tubes (Eppendorf). To confirm the successful fragmentation of chromatin, DNA from a small aliquot of each sample was purified by phenol-chloroform extraction. The size of the DNA fragments was analyzed through gel electrophoresis on a 1.6% Agarose gel with GelRed nuclei acid stain and the DNA concentration was determined by a ND-1000 spectrophotometer (Nanodrop Technologies). The rest of the chromatin was snap-frozen and stored at 80°C until further use. For chromatin immunoprecipitation, sheared chromatin was pre-cleared with BSA-blocked protein G magnetic beads (Dynabeads, Invitrogen) for 1 hour at 4°C. For every ChIP-reaction the appropriate amount of chromatin, depending on the different antibodies, was used (Supplementary Table 1). ChIP-reactions were performed by overnight incubation on a rotating wheel at 4°C in ChIP buffer (50 mM Tris-HCl at pH 8, 150 mM NaCl, 1% NP-40, 0.5% sodium deoxycholate, 20 mM EDTA, 1x Complete EDTA free protease inhibitor cocktail). As input control a 10% aliquot of each sheared chromatin sample was saved and stored at −80°C. Subsequently, 30 μl of BSA-blocked protein G magnetic beads were added to each ChIP-reaction and the mixture was incubated on a rotator at 4°C for 4 hours. The chromatin-bead complexes were washed twice at 4°C for 5 minutes at slow rotation (15 rpm) with ChIP buffer containing 0.1% SDS, twice with wash buffer (100 mM Tris-HCl pH 8, 500 mM LiCl, 1% NP-40, 1% sodium deoxycholate, 20 mM EDTA) and twice with TE buffer (10 mM Tris-HCl pH 8, 1 mM EDTA). For reverse crosslinking, chromatin-bead complexes and 10% input controls were resuspended in 300 μl elution buffer (100 mM NaHCO3, 1% SDS) with 200 mM NaCl and incubated at 65°C overnight. On the next day, the samples were treated with 1 μl of RNAse A (Thermo Scientific, EN0531) for 1 hour at 37°C. Subsequently, 6 μl 0.5M EDTA and 1 μl Proteinase K (Roche, 3115828001) were added and the samples were incubated at 45°C for 1 hour. DNA was then purified by phenol-chloroform extraction and samples were eluted in 16 μl Tris buffer (10 mM Tris-HCl pH 8) in a DNA low-binding tube and stored at −80°C.

### ChIP-sequencing

DNA concentrations for ChIP-reactions and Inputs were determined by using a Qubit dsDNA HS Assay Kit. Library construction was performed by using QIAseq Ultralow Input Library Kits (Qiagen, 180492) according to manufacturer’s protocol. The final libraries were purified by using AMPure XP beads and eluted in 23 μl nuclease free water. DNA fragment size was determined by using an Agilent 2100 Bioanalyzer and the concentration of the final libraries was measured with a Qubit dsDNA HS Assay Kit. Libraries were further validated by qPCR using a KAPA Library Quantification Kit (Roche) and pooled before being sequenced. Sequencing (single-end, 75 bp) was performed at the Genomic Core Facility (GCF, Trondheim) using an Illumina NextSeq500 sequencer with an average depth of 20 million reads per sample.

### Nuclear RNA-sequencing

Hippocampal nuclei were pooled from 3 animals per genotype. Nuclear RNA of NeuN+ and NeuN-nuclei was isolated by using the RNeasy® FFPE Kit (Qiagen, 73504) according to manufacturer’s protocol. RNA concentration and quality was determined using a Qubit RNA Assay Kit and an Agilent 2100 Bioanalyzer, respectively. Libraries were prepared by using a SMARTer® Stranded Total RNA-Seq Kit (Takara, 634485) according to the manufacturer’s instructions. Libraries were pooled and sent to BGI tech solutions, China. Sequencing (paired-end, 100 bp) was performed using aMGISEQ2000 sequencer with an average depth of 50 million reads per sample.

### Genomic data processing

#### Bilsulfite-seq data

After trimming and FastQC analysis, the clean GWBS data were aligned to the mouse reference genome GRCm38, followed by a separate extraction of genome-wide 5mC in the context of CG, CHG and CHH by using Bismark^34^ (Version 0.22.3). The 5mC count data was cleaned by filtering out low reads of coverage <4 in each cytosine. The differentially methylated regions (DMRs) were identified by HMST-Seq-Analyzer^35^. In brief, methylation regions were extracted from predefined genomic areas (e.g. TSS, TES, and gene body) DMRs predicted by robust statistical tests (Wilcoxon rank-sum test) between samples. The parameter setting was by default for gene feature regions withat least 3 mC sites in TSS and TES, 5 mC sites in gene body and the maximum distance between adjacent mC sites in a region by 200 bp.

Since HMST-Seq-Analyzer is only suited for two sample comparison: replicates of all knockouts (KO1, KO2) were compared to both replicates from WT control samples. The common DMRs among the 4 paired-wise comparison were defined as the true DMRs. The methylation difference is presented as relative ratio (rratio), which is defined by: <Methylation level in KO> – <Methylation level in WT>) /<Mean methylation of KO and WT>. DMRs were filtered by rratio > 0.1 as common hyper, and by rratio < −0.1 as common hypo. The methylation profile plots were generated by ggplot2 (version 3.4.0) using the default smoothing method of auto detection in R package. Bigwig files for genome tracks were created from bedgraphs using bedtools (v2.27.1).

#### ChIP-seq data

All datasets were quality assessed using FastQC and FastQ Screen. Fastq reads were aligned to the GRCm38/mm10 genome using Bowtie2^36^ and files were processed using samtools^37^. Visualization bigwigs for genome tracks and profile plots were generated from bam files using deepTools^38^. Correlation heatmap and profile plots were generated using deepTools and profile plot value were exported and analyzed using Graph Pad Prism 10. Peak calling was performed using epic2^39^ with an FDR cutoff of .05. Window and gap sizes were 100bp, 3 gaps for H3K4me3, 200bp and 3 gaps for H3K27me3 and 50bp, 2 gaps for Suz12. Reads were counted over consensus peaks, which were defined as peaks present in at least two of three replicates, using the Bioconductor package DiffBind (v.3.10.1) in R. Differential enrichment analysis was performed using edgeR; v.3.42.4) between genotypes and regions were annotated to the closest gene and assigned genomic feature using ChIPseeker^40^ (v1.36). Volcano plots were generated using the Bioconductor package EnhancedVolcano (v. 1.18.0) in R where differentially enriched regions (DERs) were defined as having a P value `≤` .05 and an absolute log2FC > 0.5. Pathway analysis if DERs was performed using IPA (Qiagen), pathway heatmaps were made using GraphPad Prism 10 and all Venn diagrams were made using Venny^41^. RPKM values were counted over all DERs using DiffBind and exported to Perseus^42^ where values were Z-scored and clustered using average linkage and Euclidean distance.

#### RNA-seq data

RNAseq datasets were quality assessed using FastQC and FastQ Screen (https://www.bioinformatics.babraham.ac.uk/project) before processing. Paired-end reads were aligned to the GRCm38 using Hisat2^43^ and files were processed using samtools^37^. Visualization bigwigs for genome tracks were generated from bam files using deeptools^38^. Quantification of properly aligned read pairs was done using the Rsubread package^44^ in R (v.4.3.2) and the Ensembl GRCm38 release 102 annotation. Bioconductor package edgeR^45^ (v. 3.42.4) was used to normalize counts across libraries and FPKMs were exported for plotting. Identification of differentially enriched genes (DEGs) between genotypes was performed using edgeR where DEGs were defined as having an FDR ≤ .05 and an absolute log2FC > 0.5. Volcano plots were created in R using the Bioconductor package EnhancedVolcano (v. 1.18.0) and genotypes were overlayed using Adobe Photoshop. Heatmaps were generated from FPKMs of DEGs using Perseus^42^ where rows were z-scored and hierarchical clustering was performed using average linkage and Euclidean distance. Pathway analysis of DEGs was done using Ingenuity Pathway Analysis (IPA; Qiagen) and heatmaps were made using GraphPad Prism 10. All Venn diagrams were made using Venny^41^.

#### Correlation analysis

Log2FPKM values from RNA-seq were calculated from the normalized count matrix with edgeR correcting for gene length. Transcription start sites (TSS) for all genes in the RNA count matrix were extracted from biomaRt in R and ChIP-seq reads were counted over 1000bp bins centered on the TSS using bedtools (v2.27.1). CPMs were calculated using edgeR and log2 read values were used to compute Pearson’s correlation coefficient and generate scatterplots using the R CRAN package ggpubr/ggplot2 (v.0.6/v.3.5).

#### Statistical analysis

Statistical analysis was conducted as indicated in the figure legends using GraphPad Prism software. Statistical significance was determined by Student’s t-test, one-way ANOVA or two-way ANOVA followed by appropriate post hoc test. Data represent mean ± SEM of at least three independent biological replicates. No statistical methods were used to predetermine sample size. The data distribution was assumed to be normal, but it was not formally tested.

## Results

### Loss of DNA glycosylases impairs spatial but not associative long-term memory

To explore a potential involvement of DNA glycosylases Ogg1 and Mutyh in the regulation of short-and long-term memory, we employed DNA glycosylase-deficient mice, which had previously been characterized^16^. First, we subjected 6 months old male wild type (WT), single- (Ogg1-/- or Mutyh-/-) and double-knockout (Ogg1-/-xMutyh-/-, DKO) mice to contextual fear conditioning (CFC), a hippocampus-dependent paradigm for associative learning (Figure 1A). DNA glycosylase deficient mice showed a normal response to electric foot shock exposure (Supplement Figure 1A) and similar levels of freezing during the fear-acquisition phase compared to WT (Supplement Figure 1B). Short-term and long-term memory for contextual fear, measured 1 h and 24 h, respectively after training, was not altered in DNA glycosylase deficient mice (Figure 1B i and ii) indicating that DNA glycosylases are not required for the consolidation of associative memories.

**Figure 1.**
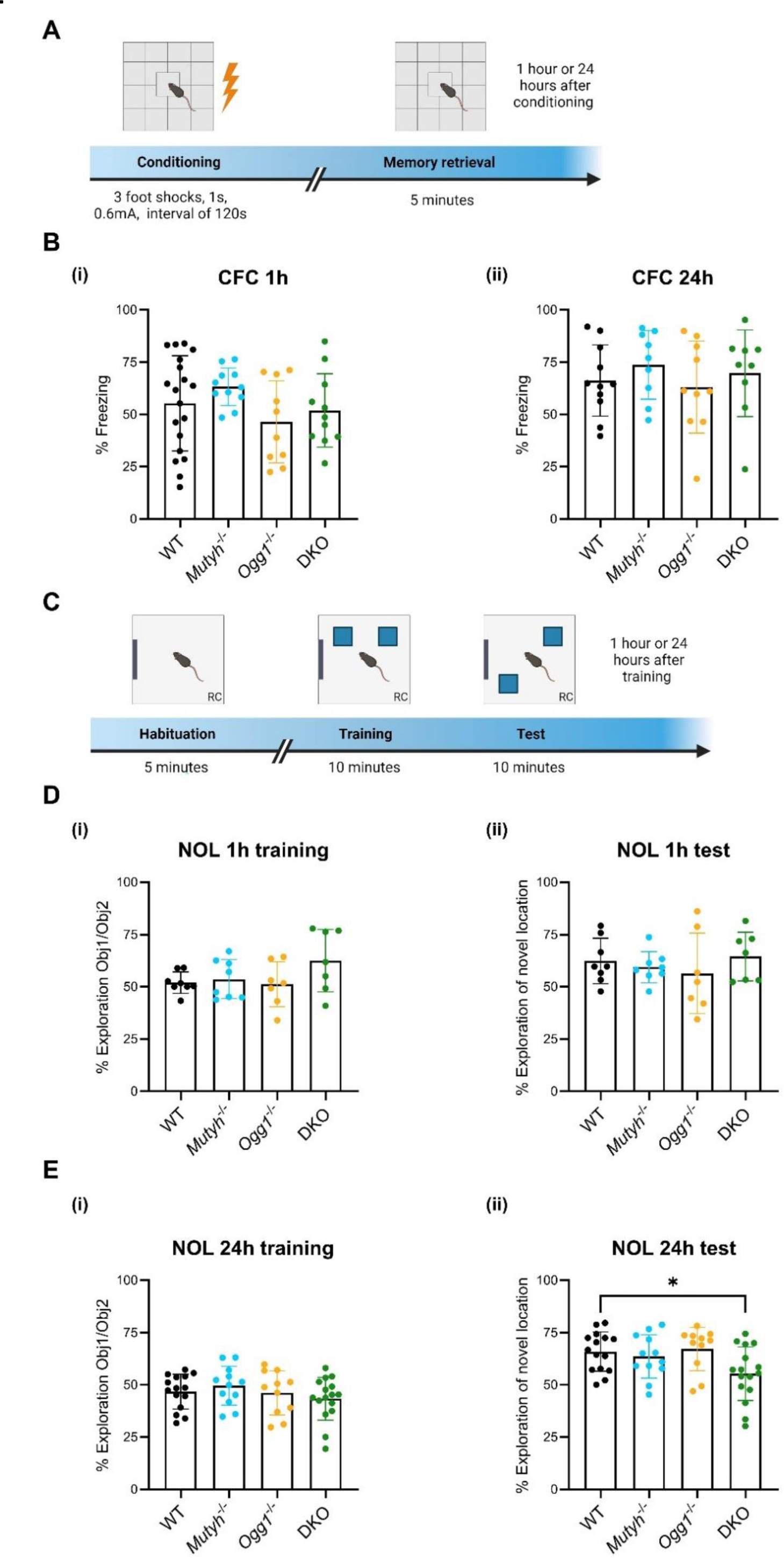
Loss of DNA glycosylases impairs spatial but not associative long-term memory. (A) Experimental design of the contextual fear conditioning (CFC) paradigm. (B) Performance in the contextual fear conditioning task measured as freezing 1 hour or 24 hours after training. At the 1 hour (i) as well as at the 24 hour (ii) training fear expression test Mutyh-/-, Ogg1-/- and DKO mice showed similar levels of freezing when compared to wild-type (WT) (WT, n = 19; Mutyh-/-, n = 11; Ogg1-/-, n = 10; DKO, n = 11). (C) Experimental design of the novel object location (NOL) memory test. (D) (i) Mutyh-/-, Ogg1-/- and DKO mice showed similar preference for the novel location and for the familiar location at the 1 hour memory retention test compared to WT (WT, n = 8; Mutyh-/-, n = 8; Ogg1-/-, n = 7; DKO, n = 7). (ii) DKO mice exhibited impaired long-term location memory by spending less time exploring the novel object at the 24 hour memory retention test compared to WT mice (WT, n = 15; Mutyh-/-, n = 12; Ogg1-/-, n = 11; DKO, n = 16, ANOVA, p = 0.0295). The relative exploration time is expressed as a percent discrimination index (D.I. = tnovel location / (tnovel location + tfamiliar location) × 100%). Data are presented as mean ± SEM, n values refer to the number of mice, *p < 0.05.

Next, we monitored WT, single-knockout and DKO mice in a novel object location (NOL) memory task (Figure 1C). All mice exhibited the same travel distance (Supplement Figure 2A) and demonstrated no significant preference for either of the objects during the training phase (Figure 1D i and Figure 1E i). At the memory retention test after 1 h DNA glycosylase deficient mice showed a similar exploration of the novel object compared to WT indicating that loss of Ogg1 and/or Mutyh does not affect short-term memory (Figure 1D ii). However, 24 hr after training DKO mice exhibited a significant decrease in the exploration of the novel object location when compared to WT (Figure 1E ii). Consistently, DKO mice showed no difference in head entries for the familiar and novel object location during the 24 hr test (Supplement Figure 2B and 2C) suggesting that both Ogg1 and Mutyh are required for long-term spatial memory that relies on spontaneous exploratory behavior.

**Figure 2.**
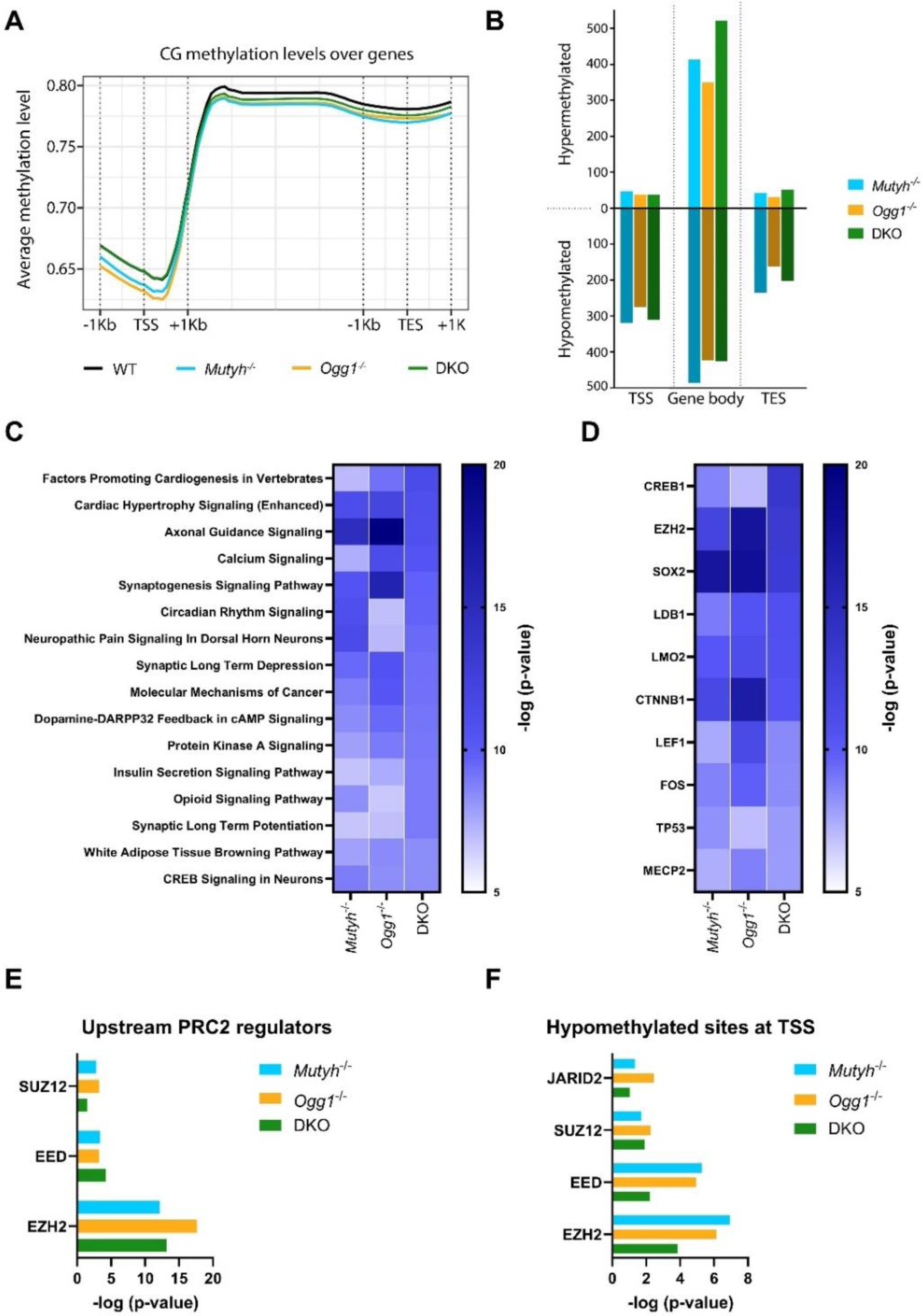
DNA glycosylases alter DNA methylation of PRC2 target genes. (A) Genome-wide average methylation level in the context of CG at transcription start site (TSS), gene body and transcription end site (TES). (B) Identification of differential methylation region (DMR) and associated genes at TSS, gene body and TES. (C) Pathways (-log10(FDR)<0.05) identified by Ingenuity pathway analysis of total DMRs. (D) Upstream regulators (-log10(FDR)<0.05) identified by Ingenuity pathway analysis of total DMRs. (E) Upstream regulators identified by Ingenuity pathway analysis of total DMRs and filtered for PRC2 subunits. (F) Chromatin immunoprecipitation Enrichment Analysis (ChEA) revealed that hypomethylated regions at TSS overlap with genes targeted by PRC2 components.

### DNA glycosylases alter DNA methylation of polycomb respressive complex 2 (PRC2) target genes

Next, we asked whether Ogg1 and Mutyh contribute to memory formation by modulating the epigenome. Epigenetic mechanisms such as DNA methylation have emerged as important regulators of memory consolidation within the nervous system^46,47^. Interestingly, several DNA glycosylases were identified as potential readers of epigenetic modifications^18,19^ and have been linked to the process of active DNA demethylation^21,48^. Therefore, we performed whole-genome bisulfite sequencing (WGBS) on hippocampal DNA from 6 months old male WT and DNA glycosylase deficient mice to map DNA methylation (5mC) at single-nucleotide resolution. Globally, mapping of 5mC to genomic locations revealed similar patterns between genotypes (Figure 2A). However, CG methylation surrounding 5’ transcription start sites (TSS) were lower in Ogg1-/- and Mutyh-/- mice, whereas at gene bodies and transcription end sites (TES) CG methylation was reduced for both single knockout and DKO mice when compared to WT.

Next, we identified differentially methylated regions (DMRs) which were either hyper- or hypomethylated at specific genomic regions between DNA glycosylase deficient mice and WT hippocampus (Figure 2B). We found 1542 regions with differential methylation in Mutyh-/-, 1279 regions in Ogg1-/-, and 1548 regions in DKO. Hypermethylated and hypomethylated regions were predominantly located within gene bodies across all genotypes. Interestingly, the majority of detected DMRs across the genomic regions were hypomethylated which was most prominent at TSS and TES in both single and double knockouts. Subsequently, we examined whether DMRs identified at TSS, gene bodies, and TES would intersect among genotypes (Supplement Figure 3). The most notable overlap of hypermethylated DMRs was detected within gene bodies across all genotypes. In contrast, we found minimal overlap of both hyper- and hypomethylated DMRs at TSS and TES among genotypes.

To understand the biological relevance and functions of altered DNA methylation we mapped all differentially methylated sites to their nearby gene annotation and performed ingenuity pathway analysis (IPA). As spatial memory impairment was found exclusively in DKO but not in single knock outs, pathways were filtered based on their significant enrichment in DKO. In the top 16 most significant pathways for single and DKO we identified an enrichment for pathways that are related to key neuronal functions including axonal guidance signaling, synaptogenesis signaling pathway, synaptic long term depression and potentiation and CREB signaling in neurons (Figure 2C).

Interestingly, enhancer of zeste homolog 2 (EZH2), a functional enzymatic component of the Polycomb Repressive Complex 2 (PRC2), was also found among the candidates with the most significant enrichment (Figure 2D). In a previous study, we discovered a significant overlap between differentially expressed genes (DEGs) identified in the hippocampus of Ogg1- and/or Mutyh-deficient mice^16^ and genes marked by H3K27me3, an epigenetic modification mediated by PRC2^49^. Thus, we examined if other subunits of PRC2 could be identified as upstream regulators across all genotypes. Indeed, we found a significant enrichment for the PRC2 subunits suppressor of zeste 12 (SUZ12) and embryonic ectoderm development (EED) in addition to EZH2 in all genotypes (Figure 2E). Furthermore, chromatin immunoprecipitation enrichment analysis (ChEA) indicated that hypomethylated regions at TSS were significantly enriched in genes targeted by PRC2 components (Figure 2F). Taken together, these results indicate that loss of Ogg1 and/or Mutyh alters DNA methylation at regulatory regions of PRC2 target genes in the adult hippocampus.

### DNA Glycosylases mediate post-translational histone modifications and Suz12 occupancy

Based on these findings, we examined whether DNA glycosylases affect the binding of the PRC2 subunit Suz12 or the deposition of post-translational histone modifications (PTMs) that are associated with PRC2 function. PRC2 is a protein complex that catalyzes the methylation of histone H3 lysine 27 (H3K27me3) leading to gene silencing and transcriptional repression^49^. In addition, PRC2 is involved in the establishment and maintenance of bivalent chromatin domains that typically feature H3K27me3, alongside the methylation of H3 lysine 4 (H3K4me3), an active mark associated with gene transcription^50^.

To decipher DNA glycosylase dependent cell-type specific epigenetic changes, we used fluorescence activated nuclear sorting (FANS) for hippocampal tissue and sorted for NeuN-positive (NeuN+) neuronal and NeuN-negative (NeuN-) non-neuronal nuclei^33^ (Figure 3A). Cell-type specificity was confirmed by the purity of the cell populations after sorting and by immunocytochemistry (ICC) post-sorting (Supplement Figure 4A and 4B). Next, we generated cell-type specific NeuN+ and NeuN-H3K4me3, H3K27me3 and Suz12 ChIP-seq data sets for all genotypes. Quality profiling of chromatin signatures for cell type-specific genes confirmed a successful enrichment of neuronal and non-neuronal nuclei. For instance, the neuronal gene Rbfox3 showed high levels of H3K4me3, an active mark, in NeuN+ nuclei, while in NeuN-nuclei H3K4me3 signals were almost completely absent and instead an enrichment for H3K27me3, a repressive mark, was found (Figure 3B). We observed no differences in the genomic distribution of H3K4me3 among genotypes in either neuronal or non-neuronal nuclei (Figure 3C). The genomic distribution of H3K27me3 occupancy at TSS within a range of plus or minus 1 kilobase (kb) was significantly higher in NeuN- for both Mutyh-/- and Ogg1-/- single knockouts. Notably, we observed a significant decrease in Suz12 occupancy at TSS in both NeuN+ and NeuN-nuclei for all genotypes when compared to WT. Annotation of the identified consensus peaks for H3K4me3, H3K27me3 and Suz12 to their genomic location revealed a similar genomic distribution across all genotypes (Supplementary Figure 5A). As expected, we identified the majority of consensus peaks for H3K4me3 to be present at promoter regions. For H3K27me3 we found that the majority of consensus peaks located in both promoter regions and distal intergenic regions in neurons for all genotypes. In glia, we observed that the majority of consensus peaks for H3K27me3 were also present in both promoter regions and distal intergenic regions, with Ogg1 and DKO showing the highest number of peaks at these locations. Suz12 consensus peaks were enriched at promoters in neuronal cells, while in NeuN- the genomic location of Suz12 seemed to be augmented in distal intergenic regions. In addition, cell-type specific allocation of consensus peaks revealed that the majority of identified peaks for H3K4me3, H3K27me3 and Suz12 were found in NeuN+ and only a subset of peaks was specific for NeuN- (Supplementary Figure 5B).

**Figure 3.**
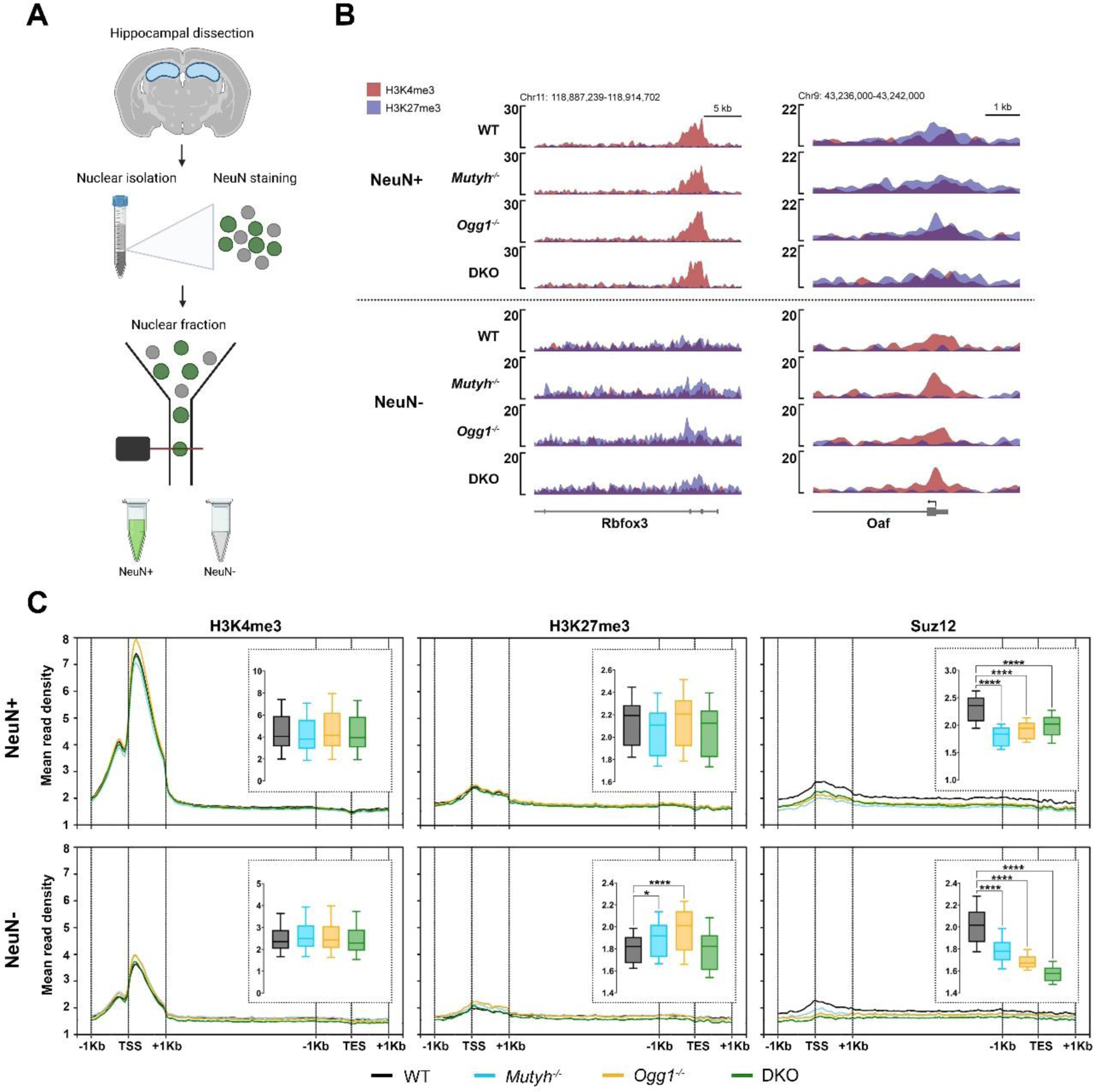
DNA glycosylases affect post-translational histone modifications and Suz12 occupancy in neuronal and non-neuronal cells. (A) Experimental design for the isolation of cell type specific nuclei used in next generation sequencing (NGS) approaches. (B) Confirmation of cell type–specific chromatin. Genome browser capture revealed an enrichment of representative H3K4me3 and H3K27me3 signals in NeuN+ and NeuN-populations at a selected neuronal gene (Rbfox3) and a glial gene (Oaf). (C) Profile plots showing mean read distribution of H3K4me3, H3K27me3 and Suz12 over transcription start site (TSS) +/- 1000bp, scaled gene body and transcription end site (TES) +/- 1000bp. Boxplots represent the distribution of mean read density over the TSS +/- 1000bp of all genes.

**Figure 4.**
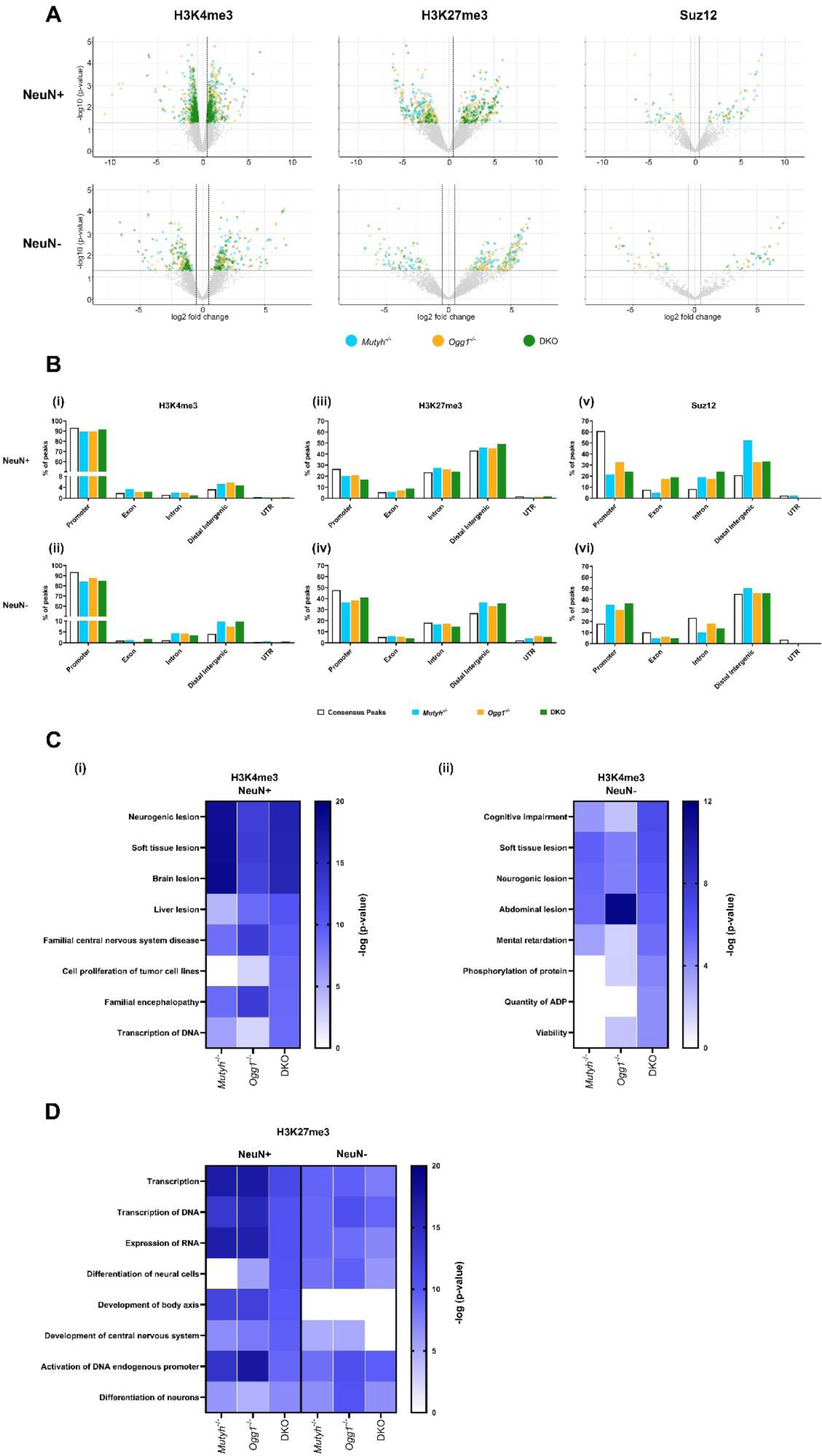
DNA Glycosylases mediate post-translational modifications associated with cognition in both neuronal and non-neuronal cells. (A) Volcano plot representation of differential enriched regions of H3K4me3 (i), H3K27me3 (ii) and Suz12 (iii) for the NeuN+ and the NeuN-population. (B) Genomic annotations of differentially enriched regions (DERs) for H3K4me3 (i), H3K27me3 (ii) and Suz12 (iii) in either the NeuN+ or NeuN-populations of Mutyh-/-, Ogg1-/- and DKO when compared to WT. (C) Disease and functions (-log10(FDR)<0.05) identified by Ingenuity pathway analysis of DERs for H3K4me3 in neurons (i) and non-neuronal cells (ii). (D) Most significant disease and functions (-log10(FDR)<0.05) identified by Ingenuity pathway analysis of DERs for H3K27me3 combined for neurons and non-neuronal cells.

### DNA Glycosylases regulate PRC2 associated PTMs linked to cognition in both neuronal and non-neuronal cells

To uncover DNA glycosylase-dependent gene specific changes in histone modification occupancy linked to PRC2 activity, we identified differentially enriched regions (DERs) in our ChIP-seq datasets. Analysis of hierarchical clustered read densities revealed a strong spatial correlation across cell types and between replicates of H3K4me3, H3K27me3 and Suz12 ChIP-assays (Supplementary Figure 6A).

The total number of DERs for both over- and under-enriched regions were similar between single-knockouts and DKO mice for H3K4me3, H3K27me3 and Suz12, respectively (Figure 4A). We identified on average 866 DERs for H3K4me3, 342 DERs for H3K27Me3 and 59 DERs for Suz12 in single-knockouts and DKO (Supplementary Figure 6B). Overall, we observed more DERs in NeuN+ nuclei compared to NeuN-nuclei across all genotypes indicating that upon loss of DNA glycosylases PTMs and Suz12 occupancy seemed to be more affected in neurons. Similar to changes in DNA methylation, we observed a minor overlap of DERs between genotypes across cell types suggesting that PTMs and Suz12 occupancy are regulated differently by Mutyh-/- and/or Ogg1-/- (Supplement Figure 7).

Next, we analyzed the genomic distribution of DERs and found that in both neuronal and non-neuronal nuclei the majority of DERs for H3K4me3 were situated in promoter regions (Figure 4B i and ii). H3K27me3 DERs showed a broader distribution and were located both at promoter regions and distal intergenic regions in NeuN+ and NeuN-nuclei (Figure 4B iii and iv). For Suz12 in neurons, we detected the highest number of DERs at distal intergenic regions across all genotypes, (Figure 4B v). In addition, we also found a high abundance of DERs at distal intergenic regions for Suz12 in non-neuronal nuclei (Figure 4B vi). Given that the majority of alterations for H3K27me3 also occur at distal intergenic regions, these findings indicate a DNA glycosylases dependent redistribution of Suz12 which consequently affects the genomic distribution of H3K27me3.

In order to comprehend the biological significance of differential H3K4me3, H3K27me3 and Suz12 occupancy, we mapped DERs to their nearby gene annotation and performed ingenuity pathway analysis (IPA) (Figure 4C and D). Since spatial memory impairment was only observed in DKO, pathways were ranked based on their significant enrichment in DKO. Among the most significant disease and function categories associated with H3K4me3 in neurons, pathways linked to various types of lesions, such as neurogenic lesions, soft tissue lesions, and brain lesions were identified for all genotypes (Figure 4C I). Moreover, we found myelination signaling pathway and axonal guidance signaling to be the top enriched canonical pathways in DKO (Supplementary Figure 8A i). In non-neuronal cells, we found an enrichment of disease and functions associated with cognition including cognitive impairment and mental retardation (Figure 4C ii). Unlike H3K4me3, the diseases and functions identified for H3K27me3 showed a strong overlap in both NeuN+ and NeuN-nuclei. Interestingly, the majority of the top identified disease and functions were related to gene regulation including transcription, transcription of DNA, expression of RNA and activation of endogenous promoter in all genotypes (Figure 4D). In non-neuronal cells we identified synaptogenesis signaling pathway and activation of NMDA receptors and postsynaptic events as the highly enriched canonical pathways for H3K27me3, most significantly in DKO (Supplementary Figure 8B ii). Because fewer DERs were identified for Suz12, representative pathway analysis could not be obtained. Our data indicate that DNA glycosylase-dependent epigenetic changes can be associated with key functions and pathways relevant for cognition in both neuronal and non-neuronal cells.

### DNA glycosylases mediate gene expression crucial for neuronal function and signaling

In a previous study, we performed bulk RNA-sequencing and have shown that Ogg1 and Mutyh regulate hippocampal gene expression related to cognition and behavior^16^. To decipher whether this is mediated in a cell type specific manner, we performed RNA-sequencing on sorted neuronal and non-neuronal nuclei. Cell-type specific RNA expression was confirmed through gene ontology analysis, which revealed neuronal processes predominantly in NeuN+ and glial-related processes in NeuN-nuclei (Supplement Figure 9A and 9B). Hierarchical clustering analysis revealed differences in gene expression levels among cell types and genotypes (Figure 5A). Next, we identified differentially expressed genes (DEGs) and found that across all genotypes DEGs were primarily downregulated in both neuronal and non-neuronal nuclei (Figure 5B). However, this was most prominent in Ogg1-/- and DKO neuronal cells with an up to 5 - 6 times higher number of downregulated genes compared to upregulated. The majority of DEGs were identified to be NeuN+ or NeuN-specific while only a minority of genes were found to be differentially expressed in both cell types, indicating that DNA glycosylases affect gene expression in a cell-type specific manner (Figure 5C).

**Figure 5.**
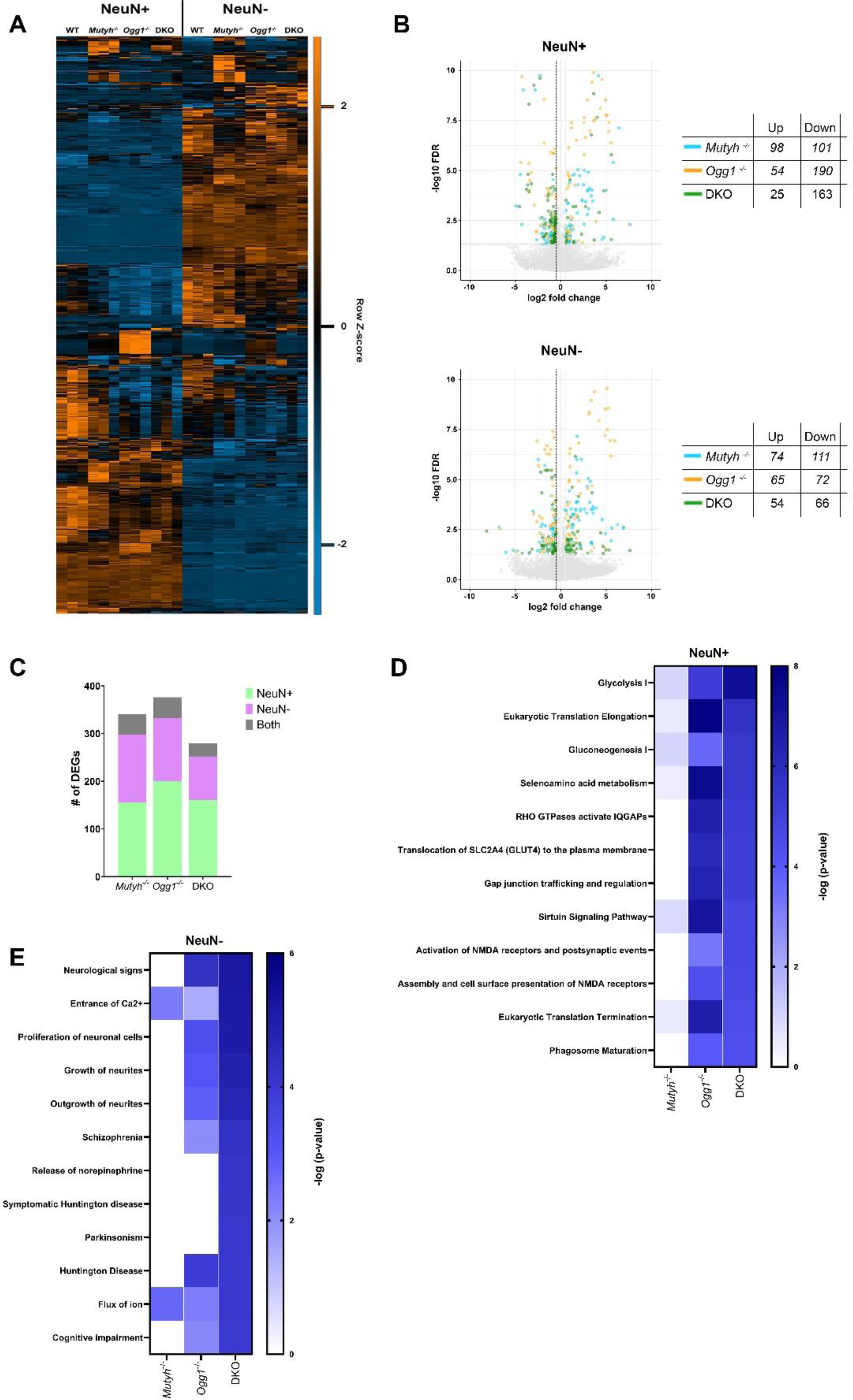
DNA glycosylases regulate gene expression important for neuronal function and signaling. (A) Heatmap over all DEGs (fold-change ≤ 0.5, FDR ≤ 0.05), clustered using Euclidean distance and average linkage. Values are FPKMs, z-scored by row across cell type showing that DEGs are predominantly found in cell type specific genes. (B) Volcano plot representation of differential expressed genes across genotypes in neuronal and non-neuronal nuclei. Tables indicate the number of genes either up- or downregulated. (C) Cell-type specific differentially expressed genes (DEGs) across Mutyh-/-, Ogg1-/- and DKO mice. Differentially expressed genes were detected exclusively in either NeuN+ or NeuN-cell populations, or in both. (D) Pathways (-log10(FDR)<0.05) identified by Ingenuity pathway analysis of DEGs in neurons. (E) Pathways (-log10(FDR)<0.05) identified by Ingenuity pathway analysis of DEGs in non-neuronal cells.

Next, we examined whether DEGs would intersect among genotypes (Supplement Figure 10). The most pronounced overlap was observed in the downregulated DEGs in neurons across all genotypes. Although we did not find a strong overlap of DEGs across all genotypes, we found an overlap of DEGs between Ogg1-/- and DKO of downregulated DEGs in neuronal and non-neuronal nuclei.

Next, we examined the biological implications of DEGs and observed a significant enrichment of several metabolic pathways, including glycolysis I, gluconeogenesis I, and selenoamino acid metabolism, with a particularly pronounced enrichment in Ogg1-/- and DKO within neurons (Figure 5D). Interestingly, we also discovered pathways associated with NMDA receptors, with the strongest enrichment in DKO. In glial, pathway analysis revealed various diseases and functions including entrance of Ca2+, neurite growth, schizophrenia, Huntington’s disease, Parkinsonism, and cognitive impairment, most significantly enriched in DKO (Figure 5E). In summary, although we identified cell-type specific changes in gene expression upon loss of DNA glycosylases, DEGs in both cell-types were associated with highly relevant categories for neuronal function and cognition, particularly in DKO.

### Post-translational histone marks modulated by DNA glycosylases regulate gene expression

Next, we explored a potential correlation between DNA glycosylase-dependent epigenetic and transcriptional changes (Figure 6A). Spearman correlation analysis between FPKMs (fragments per kilobase million) obtained through RNA-seq and CPMs (counts per million) obtained from ChIP-seq revealed that the correlation of PTMs (H3K4me3 and H3K27me3) was consistent across all genotypes. DNA glycosylase dependent changes of H3K4me3 CPMs and RNA FPKMs were both positively correlated in neurons and non-neuronal cells. In contrast, H3K27me3, which is associated with transcriptional repression, demonstrated a negative correlation with RNA changes. In addition, we observed a minor negative correlation between Suz12 occupancy and RNA expression. To examine a potential correlation between DNA glycosylase-dependent DNA methylation and gene expression changes we intersected DMRs with DEGs. Venn diagrams revealed only a minor overlap of DEGs and DMRs across all genotypes (Supplement Figure 12).

**Figure 6.**
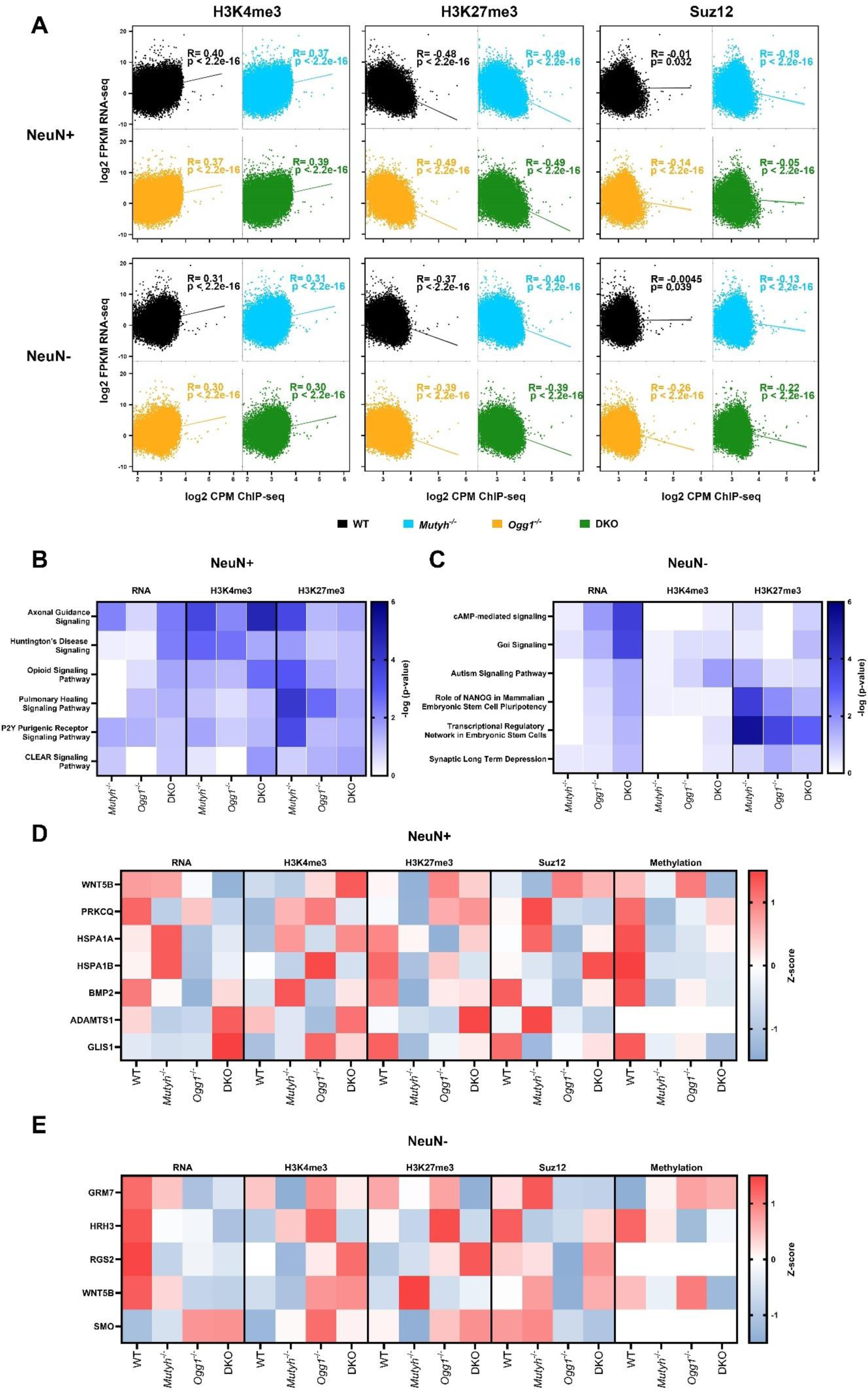
Post-translational histone marks regulated by DNA glycosylases influence gene expression. (A) Spearman correlation analysis between ChIP-seq and RNA-seq data. Scatterplot showing the correlation of H3K4me3, H3K27me3, and Suz12 CPMs (counts per million) in the transcription start site (+/- 500bp) with RNA FPKMs (fragment per kilobase million) for neurons and non-neuronal cells. (B) Overlapping pathways for RNA, H3K3me3 and H3K27me3 (-log10(FDR)<0.05) identified by Ingenuity pathway analysis in neurons. (C) Overlapping pathways for RNA, H3K3me3 and H3K27me3 (-log10(FDR)<0.05) identified by Ingenuity pathway analysis in non-neuronal cells. (D) Differences in gene expression (FPKMs), DNA methylation (mean signal) and occupancy for post translational histone modifications (PTMs) and Suz12 (CPMs in the TSS) for overlapping PRC2 target genes associated with axonal guidance, Huntington’s disease and opioid signaling in neurons. (E) Differences in gene expression (FPKMs), DNA methylation (mean signal) and occupancy for PTMs and Suz12 (CPMs in the TSS) for overlapping PRC2 target genes associated with cAMP-mediated signaling, gαi signaling and Autism signaling pathway in non-neuronal cells

Since we found only minimal gene-specific overlap between DMRs, DERs and DEGs, we wondered whether similar biological processes are affected. Therefore, we intersected the pathways identified in each individual dataset that showed significant enrichment. Interestingly, in neurons the overlap uncovered numerous pathways linked to critical neuronal processes and neurodegenerative diseases such as axonal guidance signaling, opioid signaling pathway and Huntington’s disease. Notably, DKO exhibited the strongest enrichment in the axonal guidance signaling pathway for both RNA and H3K4me3 (Figure 6B). In glia, overlapping pathways were less prominent but we found key neuronal processes and diseases including cAMP-mediated signaling, synaptic long term depression and Autism signaling pathway, all with most significant enrichment in DKO for RNA (Figure 6C).

We then compared RNA expression (FPKMs), DNA methylation, Suz12 occupancy, and post-translational modifications (PTMs) of genes identified in the overlapping pathways involved in key neuronal processes across all genotypes (Figure 6D and E). Importantly, all the presented genes, are known to be target genes for PRC2. Interestingly, in DKO we found a negative correlation between RNA and H3K27me3 for several genes (in neurons: WNT5B, PRKCQ, HSPA1A; in glia: RGS2). Additionally, ADAMTS1 and GLIS1 showed a correlation between RNA and H3K4me3 in neurons for DKO. Together, our findings suggest that the DNA glycosylases Ogg1 and Mutyh affect gene expression important for neuronal functions and signaling by modulating the epigenetic landscape through post-translational histone modifications and DNA methylation

## Discussion

While DNA glycosylases are conventionally associated with preserving genome integrity, our findings establish Ogg1 and Mutyh as important regulators of the neuronal epigenome, essential for spatial memory formation in the hippocampus. Our study demonstrates that in the hippocampus, Ogg1 and Mutyh play a role in regulating DNA methylation, particularly affecting the methylation status of PRC2 target genes. Moreover, we find that Ogg1 and Mutyh modulate histone modifications associated with PRC2 activity in neuronal- and glial-specific nuclei of the hippocampus. Our results indicate a novel role for Ogg1 and Mutyh beyond DNA repair in modulating the neuronal epigenome to regulate transcriptional responses important for neuronal function and signaling.

### DNA glycosylases regulate spatial long-term memory

Both our research and other studies have highlighted the crucial involvement of DNA glycosylases in higher brain functions associated with cognition^51,52^. In a previous study, we evaluated the learning and memory performance of single and double knockout mice deficient for Ogg1 and Mutyh by conducting a Morris water maze test. DKO mice were significantly slower in learning the position of the platform compared to wild-type mice but showed no difference in memory retention leaving the question if both Ogg1 and Mutyh affect memory formation unanswered^16^. Here, we demonstrate that the absence of Ogg1 and/or Mutyh does not influence short-term and long-term memory for contextual fear, suggesting that Ogg1 and Mutyh are not essential for the consolidation of associative memories. In a novel object location memory task assessing spontaneous exploratory behavior, DKO mice exhibited no impairment in short-term memory but displayed a deficit in long-term spatial memory. Interestingly, we did not detect any memory formation impairment in Ogg1 and Mutyh single knockouts. Only the deletion of both DNA glycosylases led to a deficit in memory retention, indicating that memory formation is regulated synergistically by both DNA glycosylases. Our findings suggest that the presence of both Ogg1 and Mutyh is particularly important in memory tasks that recruit an innate behavioral preference for novelty and do not involve exogenous reinforcers such as swim stress or an electric foot shock. Our results are consistent with other studies^53,54^ demonstrating that stressful reinforcement stimuli might trigger alternative mechanisms that could compensate for the absence of both Ogg1 and Mutyh in our mouse model.

### DNA glycosylases modulate DNA methylation and PTMs associated with PRC2 function

Oxidative DNA damage and its repair has been demonstrated to be involved in transcriptional regulation but whether these processes are directly linked has remained largely unexplored. We have previously shown that loss of Ogg1 and/or Mutyh impact gene transcription independent of global accumulation of 8-oxoG in the hippocampus^16^. However, the underlying mechanism how DNA glycosylases mediate gene expression independent of DNA repair remain elusive.

Here, we uncover a novel function for Ogg1 and Mutyh, extending beyond their conventional role in DNA repair, by modulating DNA methylation in the adult mouse hippocampus. DMRs across all genotypes were predominantly hypomethylated and at TSS the majority of hypomethylated DMRs were significantly enriched in genes targeted by PRC2 in both single and double knockouts. Consistent with our findings, increasing evidence indicates that DNA glycosylases play a role in modulating cellular function by influencing epigenetic alterations. Ogg1 and Mutyh have been demonstrated to bind to, but not to excise, 5mC, potentially preventing DNA demethylation^19,55^. Interestingly, it has been shown that Ogg1 is part of a multiprotein complex that includes DNMT1 and EZH2, which facilitates CpG promoter methylation and induces alterations in post-translational histone modifications such as H3K4me3 and H3K27me3^22^. Consistently, we observed that the absence of Ogg1 and Mutyh disrupts the occupancy of both marks H3K4me3 and H3K27me3. The precise mechanism by which PRC2 is recruited to its target sites to promote transcriptional silencing remains unclear. We suggest that Ogg1 and Mutyh mediate the recruitment of PRC2 to its target genes. In line with this, we found a redistribution of H3K27me3 and Suz12 upon loss of Ogg1 and Mutyh indicating that PRC2 mediated deposition of H3K27me3 occurs at random genomic locations and not at its target genes. We believe that recruitment of PRC2 by Ogg1 and Mutyh is unlikely to occur through 8-oxoG. Mutyh recognizes adenine opposite 8-oxoG which is incorporated after replication^56^. However, neurons are postmitotic, thus Mutyh is not needed for 8-oxoG repair. In a recent study, we profiled genome-wide distribution of 8-oxoG and found no correlation with gene expression in cells deficient for Ogg1 and Mutyh^28^. This suggests that 8-oxoG and DNA Glycosylases modulate gene transcription independently arguing against an 8-oxoG-dependent mechanism. Alternatively, Ogg1 and Mutyh may recruit PRC2 to its targets sites by recognizing secondary DNA structures such as G-quadruplex (G4) structures. G4 structures occur at CpG rich regions and have been implicated in gene regulation^57,58^. Although the potential association between DNA glycosylases and G4 structures has been suggested^28,59^, further research is required to elucidate this potential mechanism.

### DNA glycosylases regulate the expression of genes targeted by PRC2 important for neuronal function

In the adult brain, PRC2 plays a crucial role in maintaining neuronal function and survival by transcriptionally silencing specific bivalent genes. Interestingly, mice lacking PRC2 in neurons showed signs of progressive neurodegeneration, with alterations in gene expression linked to Huntington’s disease^60^. In addition, it has been demonstrated that PRC2 is crucial for the activation of microglia and that loss of PRC2 causes changes in neuronal morphology and complex behaviors associated with neurodegenerative diseases^61^. Consistently, we observed in previous studies that upon loss of Ogg1 and Mutyh DEGs exhibited the highest enrichment in target genes of PRC2 in a cancer cell line^28^ and a significant overlap with genes harboring the epigenetic mark H3K27me3 in the mouse hippocampus^16^. In line with these finding, we found that upon loss of Ogg1 and Mutyh cell-type specific alterations in gene expression were associated with highly relevant categories for neuronal function and cognition in both neurons and non-neuronal cells. Moreover, in glia, lack of Ogg1 and Mutyh led to gene expression changes associated with Huntington’s disease, Parkinsonism and cognitive impairment. In addition, several genes exhibiting a significant enrichment within those pathways have been reported to be PRC2 target genes. Together, our findings indicate that Ogg1 and Mutyh may cooperate with PRC2 in modulating the epigenome and consequently, gene transcription.

In conclusion, our findings reveal a novel function for Ogg1 and Mutyh beyond DNA repair, as modulators of the epigenome to regulate transcriptional responses crucial for neuronal function and signaling. Notably, we observed the most pronounced effects in DKO, which aligns with our finding that only the simultaneous deletion of Ogg1 and Mutyh results in impaired memory formation. We found alterations in DNA methylation, occupancy of PTMs and Suz12, and RNA expression in both single-knockouts and DKO, with minimal overlap across genotypes. This implies that the behavioral phenotype arises specifically from the combined deficiency of Ogg1 and Mutyh. Consequently, gene expression underlying memory formation appears to be regulated synergistically by both DNA glycosylases. In future studies, it will be interesting to examine the role of Ogg1 and Mutyh in epigenetic maintenance underlying neurodegenerative diseases.

## Acknowledgements

We thank the Genomic Core Facility (GCF) at NTNU for the help to obtain high-throughput sequencing data.

