## Supplementary Data for "DNA glycosylases Ogg1 and Mutyh mediate gene expression of PRC2 target genes important for neuronal processes underlying memory formation"

**Supplementary table 1**

**Supplementary figures 1- 12**

| Antibody | Raised in | Company | # | Chromatin Input | Ab used per IP reaction |
|---|---|---|---|---|---|
| H3K27trime | mouse | Abcam | ab6002 | 2 µg | 5 µg |
| H3K4trime | rabbit | Abcam | ab8580 | 2 µg | 5 µg |
| Suz12 | rabbit | Cell Signaling | #3737S | 4.5 µg | 1 µg |

**Supplementary Table 1. Antibody list and chromatin input for chromatin Immunoprecipitation**

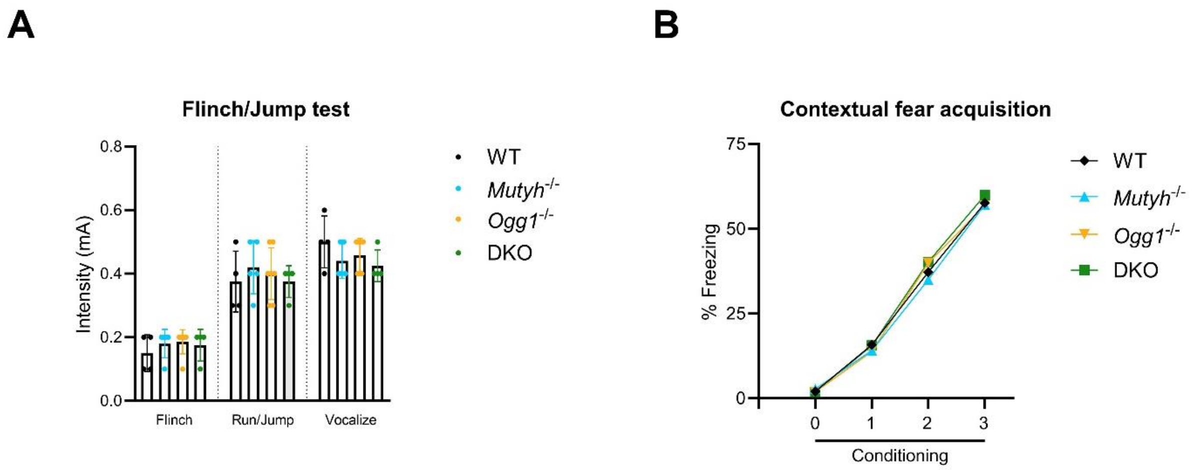


**Supplementary figure 1. DNA glycosylase deficient mice show no abnormalities in the contextual fear paradigm**

(A) DNA glycosylase deficient mice show a normal response to electric foot shock. Flinch/Jump test revealed similar general responses to electric foot shock between Mutyh-/-, Ogg1-/- and DKO mice when compared to WT mice (WT, n = 4; Mutyh-/-, n = 5; Ogg1-/-, n = 7; DKO, n = 4). Mice were scored based on their first visible response to the foot shock (flinch), their first pronounced motor response (run or jump), and their first vocalized distress as described in Materials and Methods. Data are presented as mean ± SEM, n values refer to the number of mice.

(B) In the contextual fear conditioning paradigm, DNA glycosylase deficient mice showed similar levels of freezing to WT during the fear-acquisition phase. Average freezing presented per genotype throughout conditioning (0=acclimatisation, 1,2,3=sequential electric foot shocks; WT, n = 11; Mutyh-/-, n = 9; Ogg1-/-, n = 10; DKO, n = 9; n values refer to the number of mice).

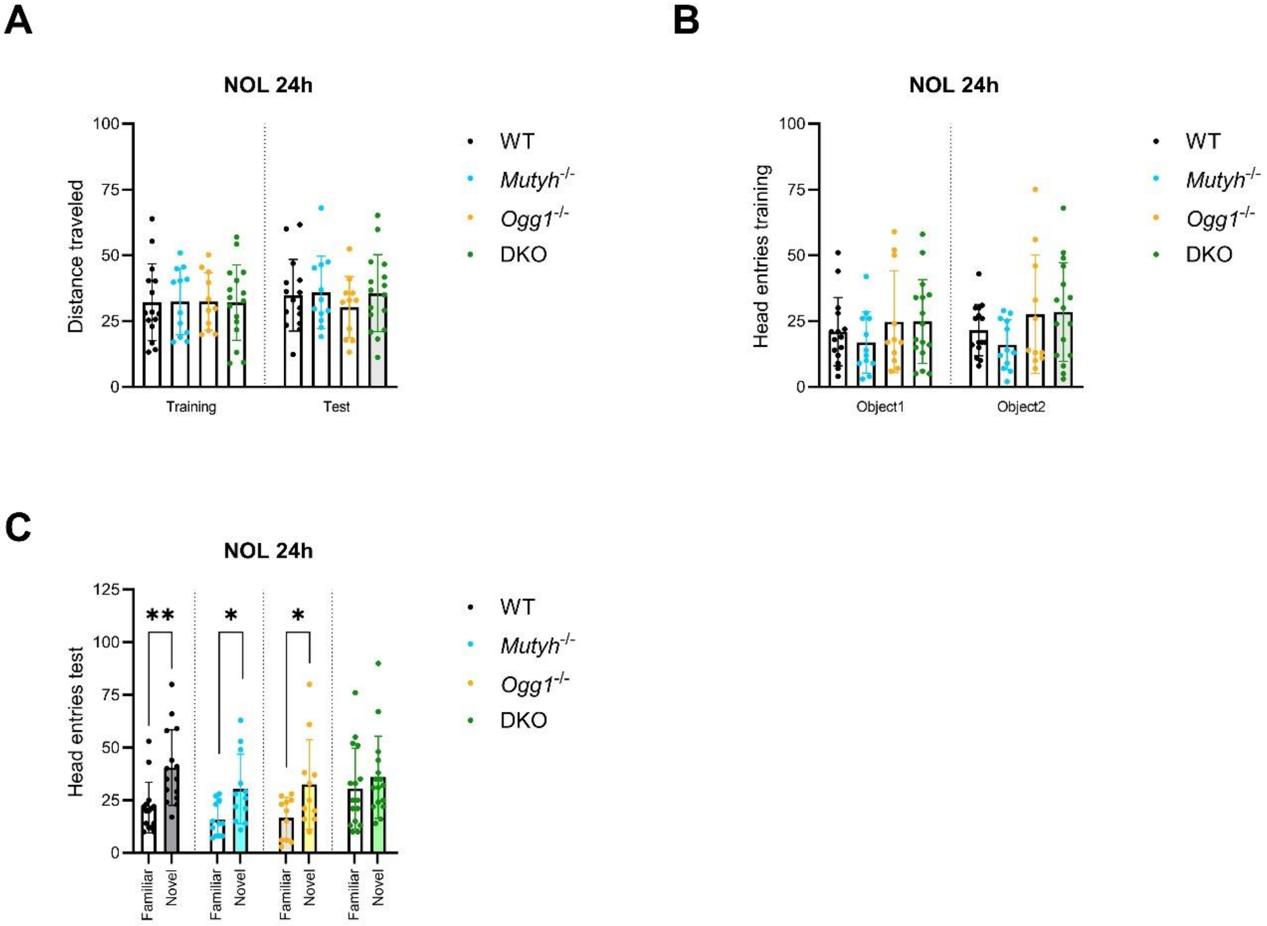


**Supplementary figure 2. Novel object location (NOL) paradigm revealed a difference in head entries for DKO mice during the 24 hour NOL test**

(A) Analysis of the distance traveled (in meters) during the novel object location paradigm revealed no difference between WT, Mutyh-/-, Ogg1-/- and DKO mice during training and the 24 hour test (WT, n = 15; Mutyh-/-, n = 12; Ogg1-/-, n = 11; DKO, n = 16; n values refer to the number of mice).

(B) Head entries examined during training indicated that DNA glycosylase deficient and WT mice (n = 15) had no preference for either of the objects (Mutyh-/-, n = 12; Ogg1-/-, n = 11; DKO, n = 16).

(C) During the NOL test for 24 hours WT, Mutyh-/-, Ogg1-/- showed a significant increase in head entries for the novel object. Head entries examined for DKO mice revealed no difference between the familiar and the novel object (Student’s t-test, WT, n = 15, p = 0.002; Mutyh-/-, n = 12, p = 0.0114; Ogg1-/-, n = 11, p = 0.0355; DKO, n = 16, p = 0.4379). Data are presented as mean ± SEM, n values refer to the number of mice, *p < 0.05.

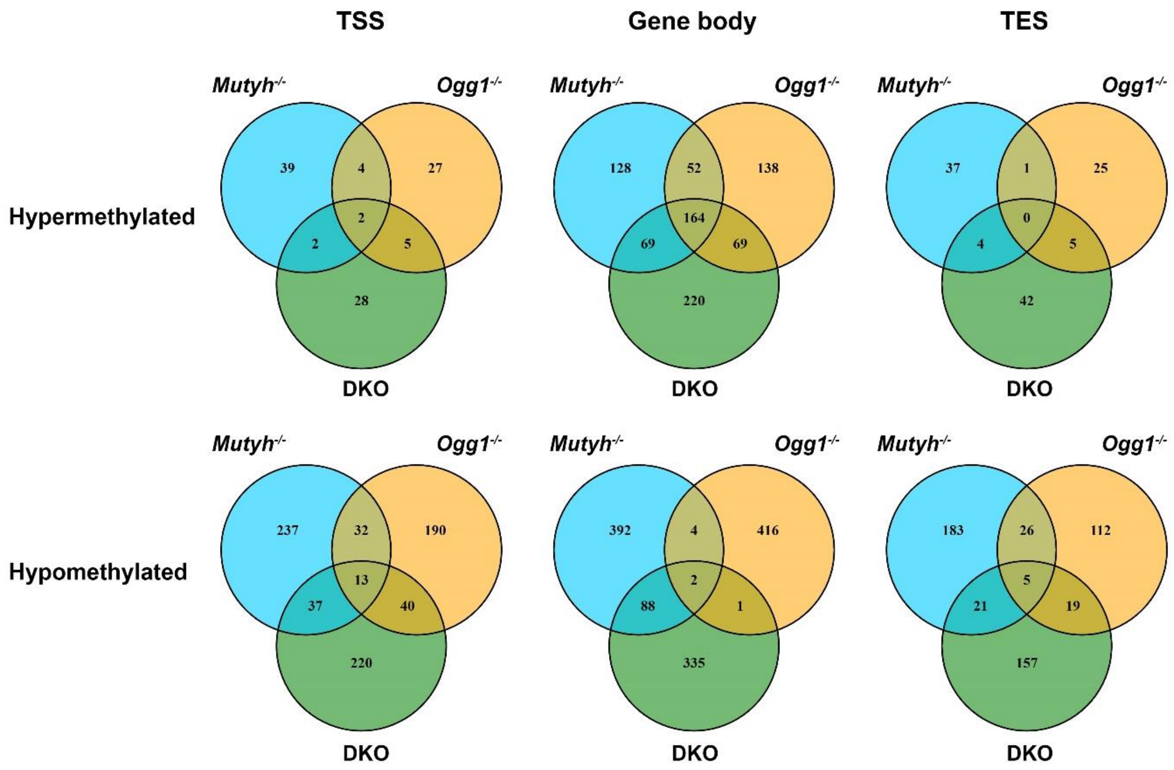


**Supplementary figure 3. Overlap of DNA Methylation across genotypes**

Venn diagrams depict the overlap of DNA methylation among genotypes for either hyper- or hypomethylated sites at the transcription start site (TSS), gene body and transcription end site (TES).

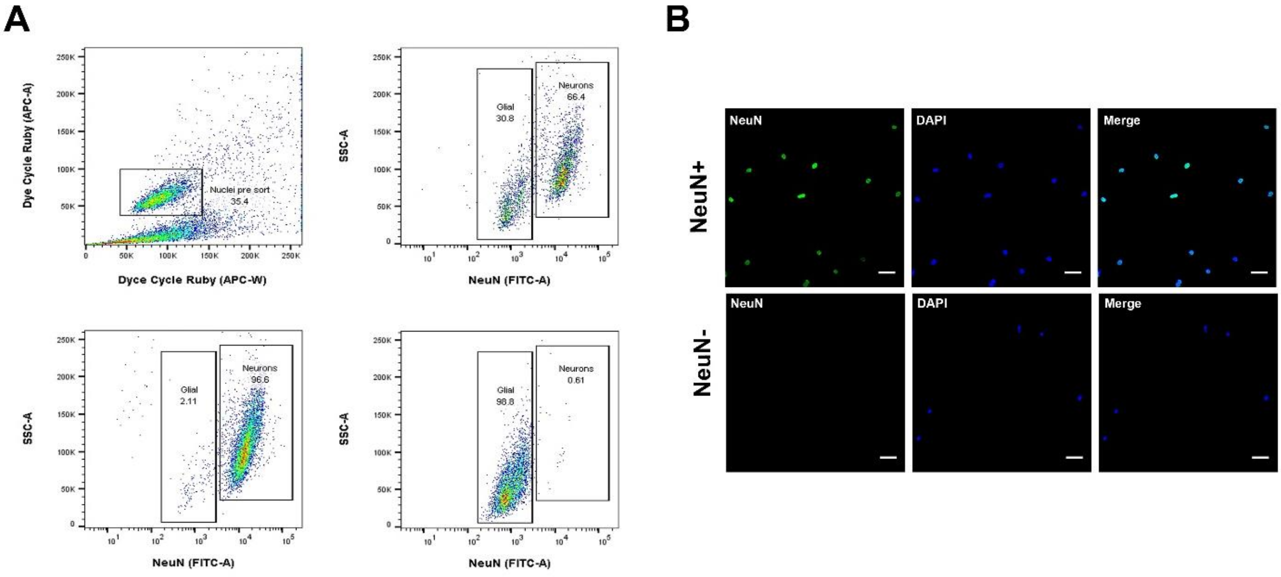


**Supplementary figure 4. Fluorescence activated nuclear sorting (FANS) of cell type-specific nuclei**

(A) FANS profiles for Dye cycle Ruby and NeuN demonstrating the gating strategy during the sorting (top). Purity of the sorting process was confirmed by probing the sorted cell populations after sorting (bottom). Percentage of the filtered population are indicated within the channel. Similar results were obtained for all three independent experiments.

(B) Confocal images of nuclei to confirm the purity of NeuN+ and NeuN- nuclei after sorting (Scale bar = 30μM).

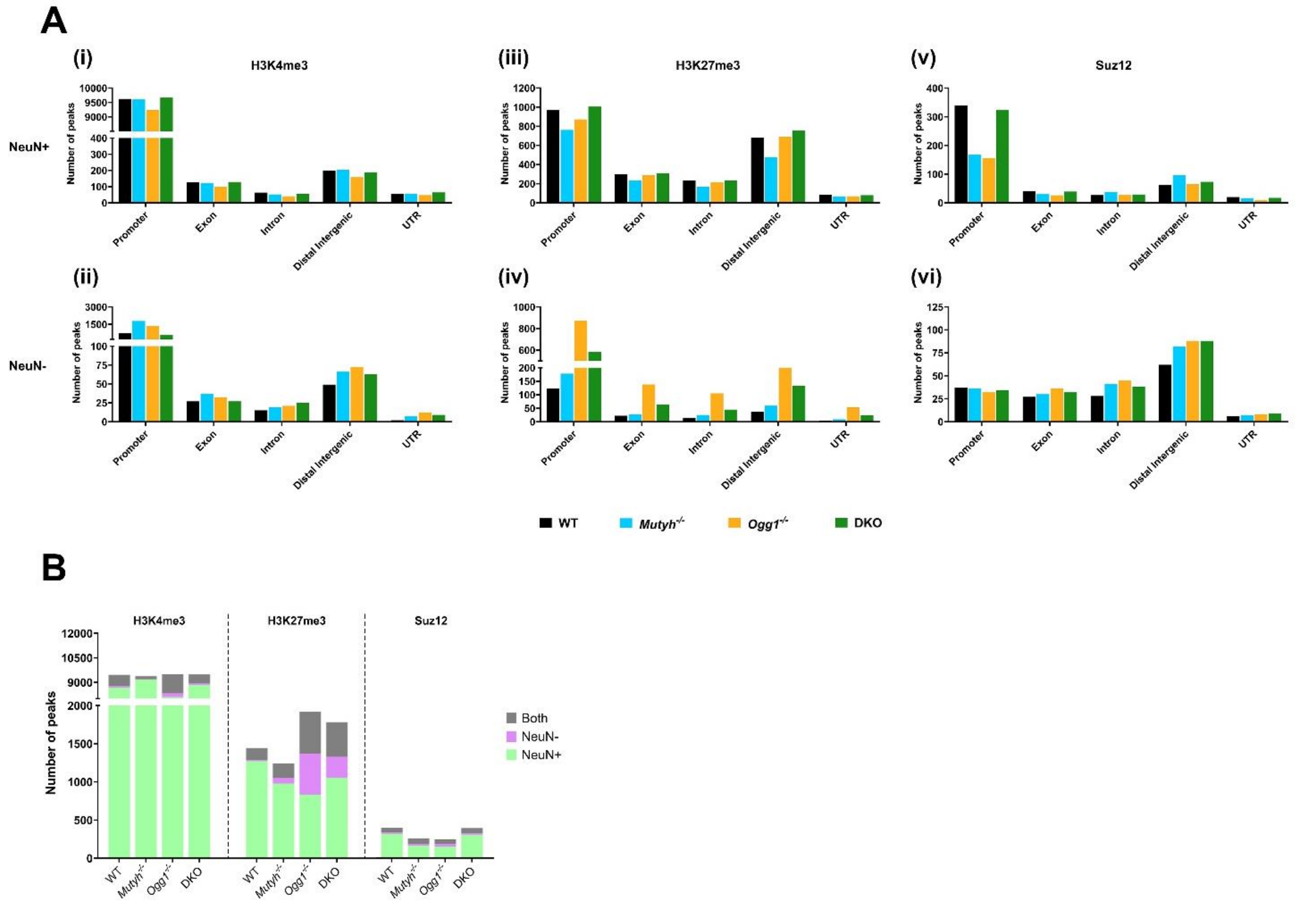


**Supplementary figure 5. Characterization of identified consensus peaks**

(A) Genomic annotations of consensus peaks for H3K4me3 (i), H3K27me3 (ii) and Suz12 (iii) in either the NeuN+ or NeuN- populations of WT, Mutyh-/-, Ogg1-/- and DKO.

(B) Cell-type specific distribution of consensus peaks for H3K4me3, H3K27me3 and Suz12 across the hippocampus of WT, Mutyh-/-, Ogg1-/- and DKO mice.

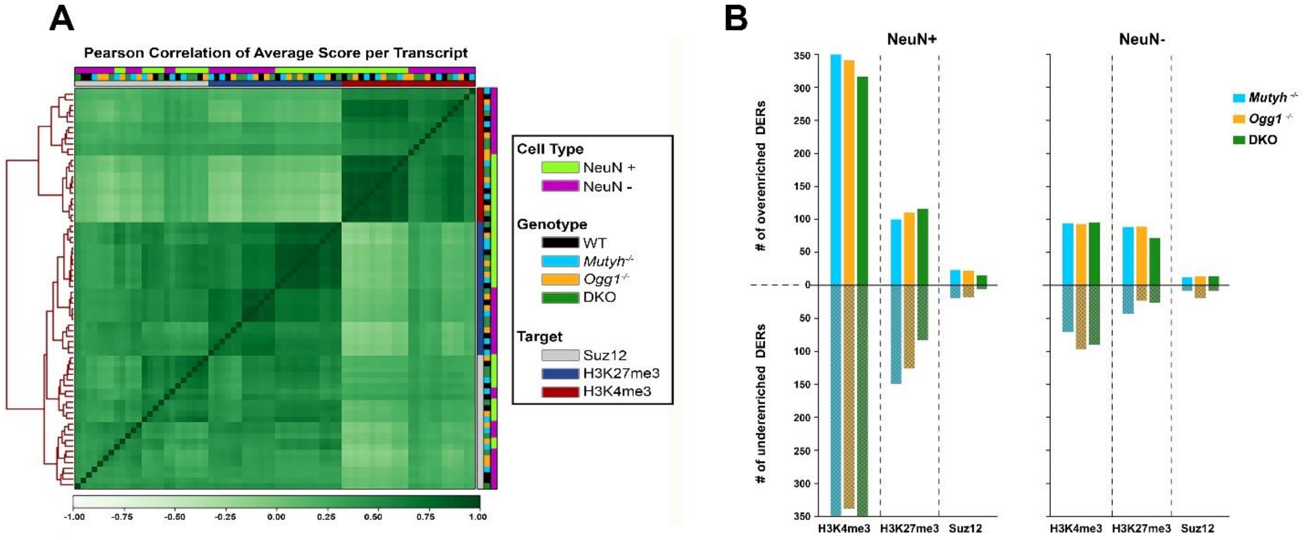


**Supplementary figure 6. Characterization of H3K4me3, H3K27me3 and Suz12 occupancy**

(A) Correlation heatmap with hierarchical clustering of read densities for H3K4me3, H3K27me3 and Suz12 replicates for NeuN+ and NeuN-.

(B) Number of differentially enriched regions for H3K4me3, H3K27me3 and Suz12 either over-enriched or under-enriched for NeuN+ and NeuN-.

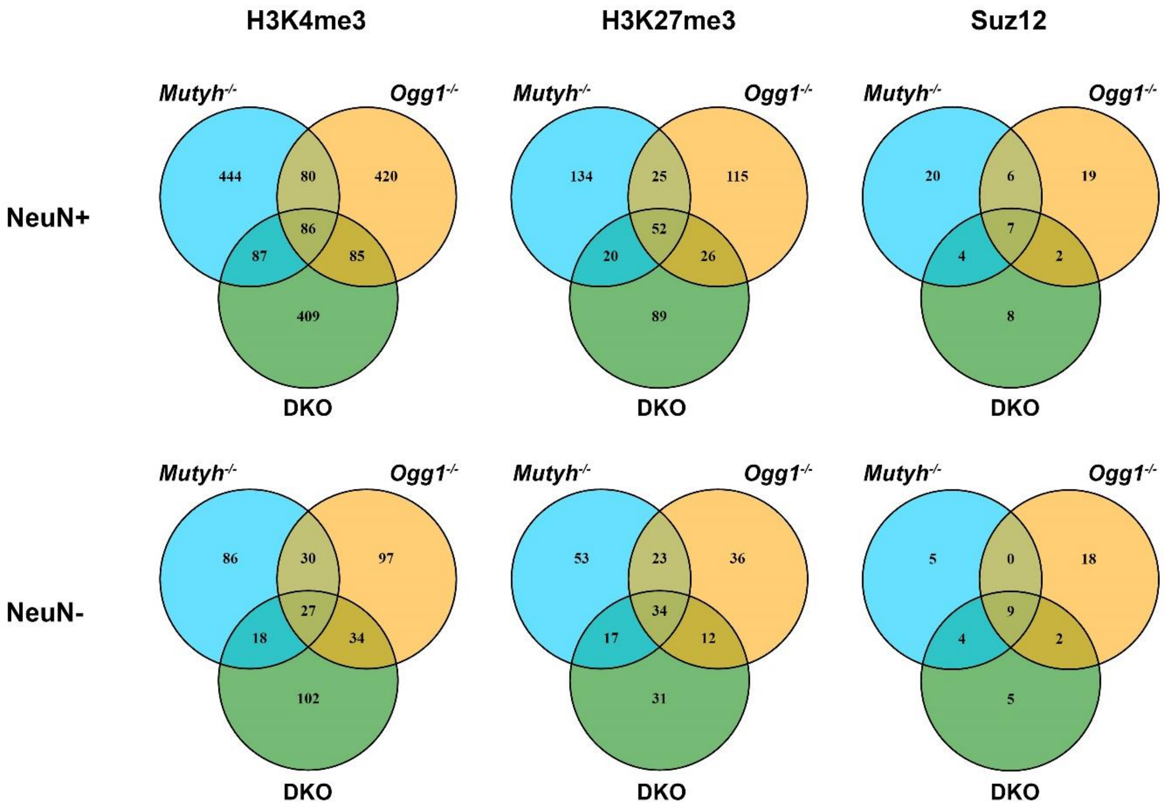


**Supplementary figure 7. Overlap of H3K4me3, H3K27me3 and Suz12 occupancy across genotypes**

Venn diagrams illustrate the intersection of H3K4me3, H3K27me3, and Suz12 occupancy across genotypes for both neurons and non-neuronal cells.

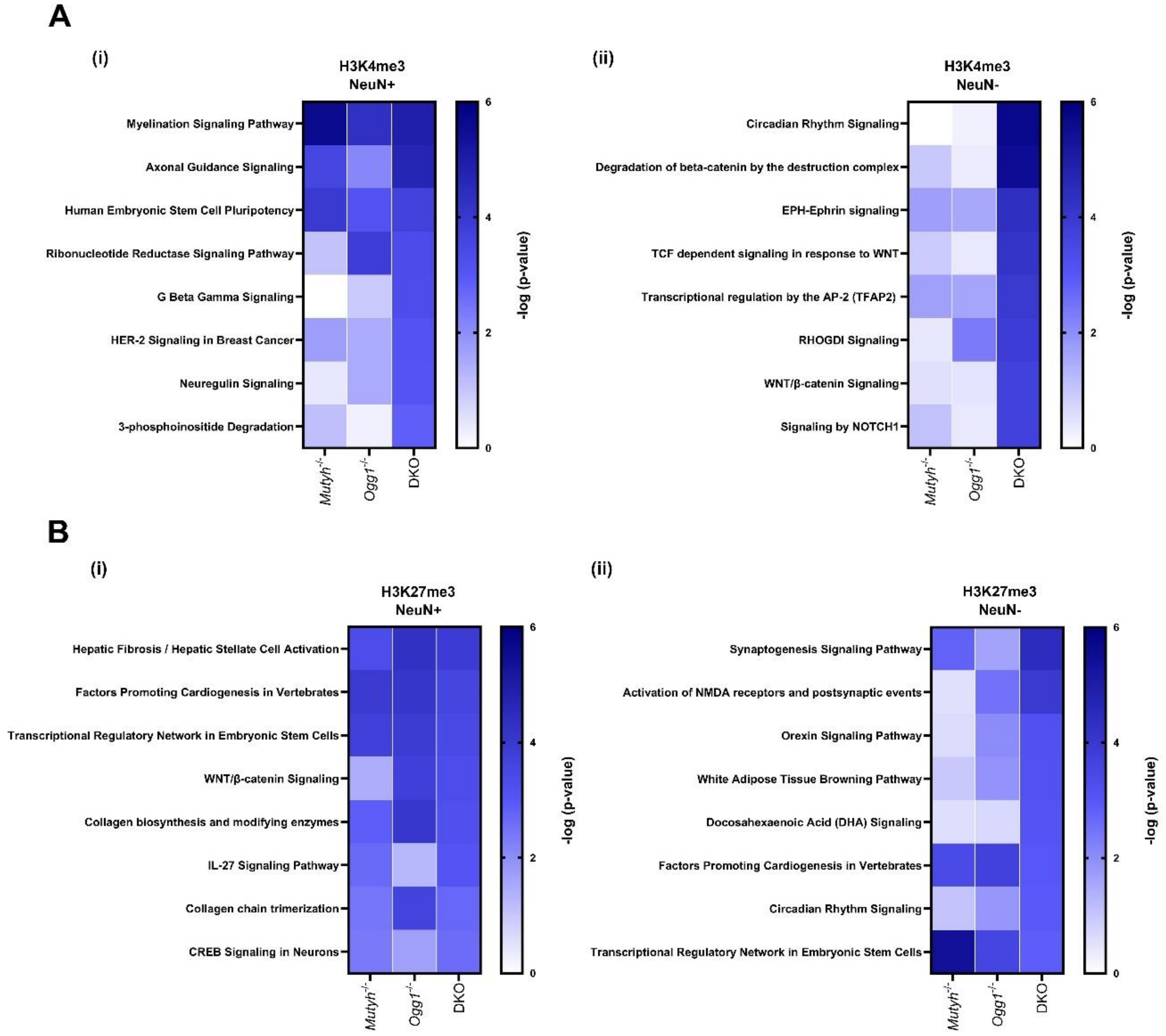


**Supplementary figure 8. Pathways for differentially enriched regions for H3K4me3 and H3K27me3**

(A) Pathways (-log10(FDR)<0.05) identified by Ingenuity pathway analysis of DERs in neurons.

(B) Pathways (-log10(FDR)<0.05) identified by Ingenuity pathway analysis of DERs in non-neuronal cells.

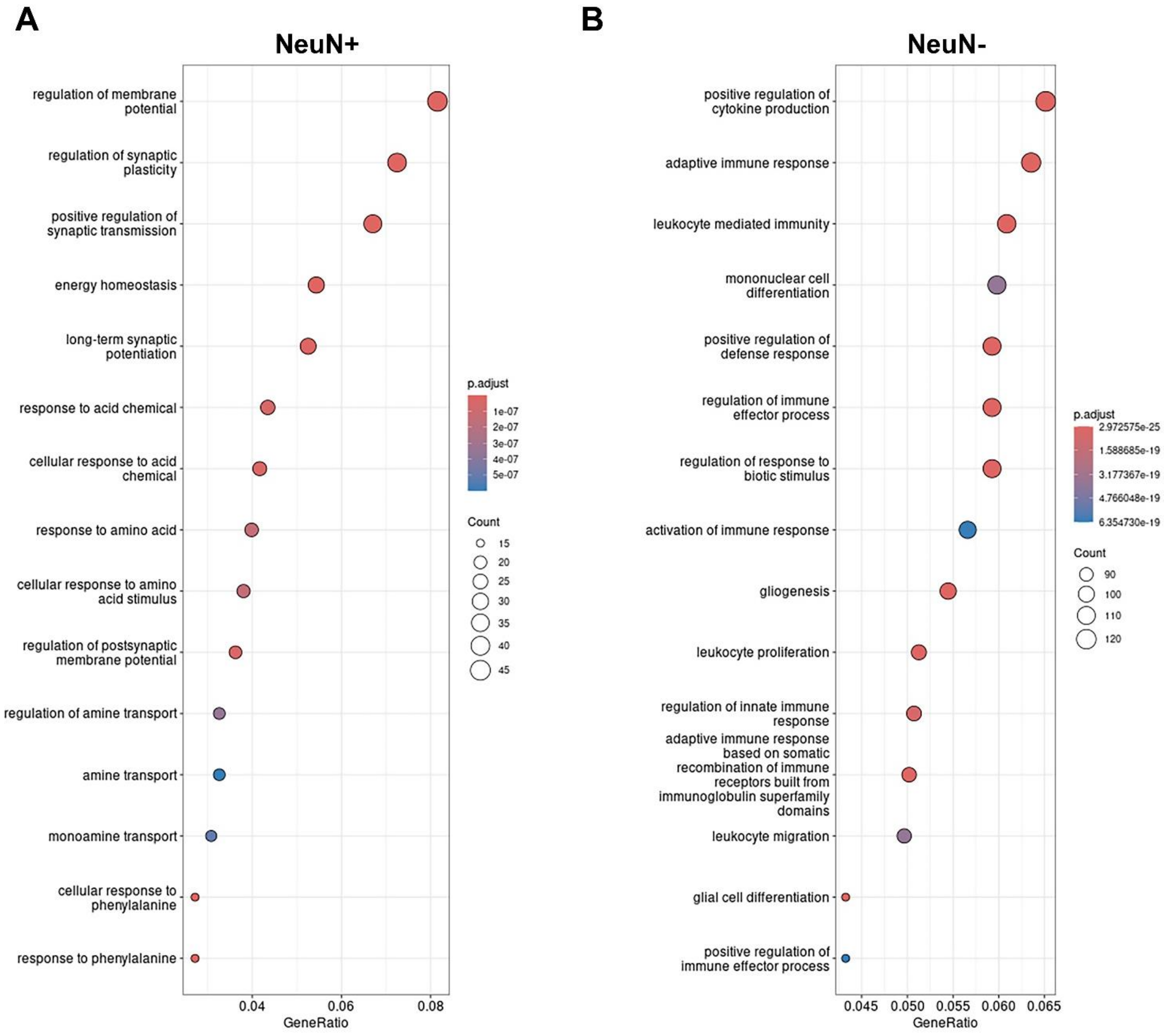


**Supplementary figure 9. Isolation of cell-type specific nuclei by fluorescence activated nuclear sorting (FANS) for RNA-sequencing**

Gene ontology analysis revealed cell-type specific RNA expression in neurons (A) and non-neuronal cells (B) in wild type mice.

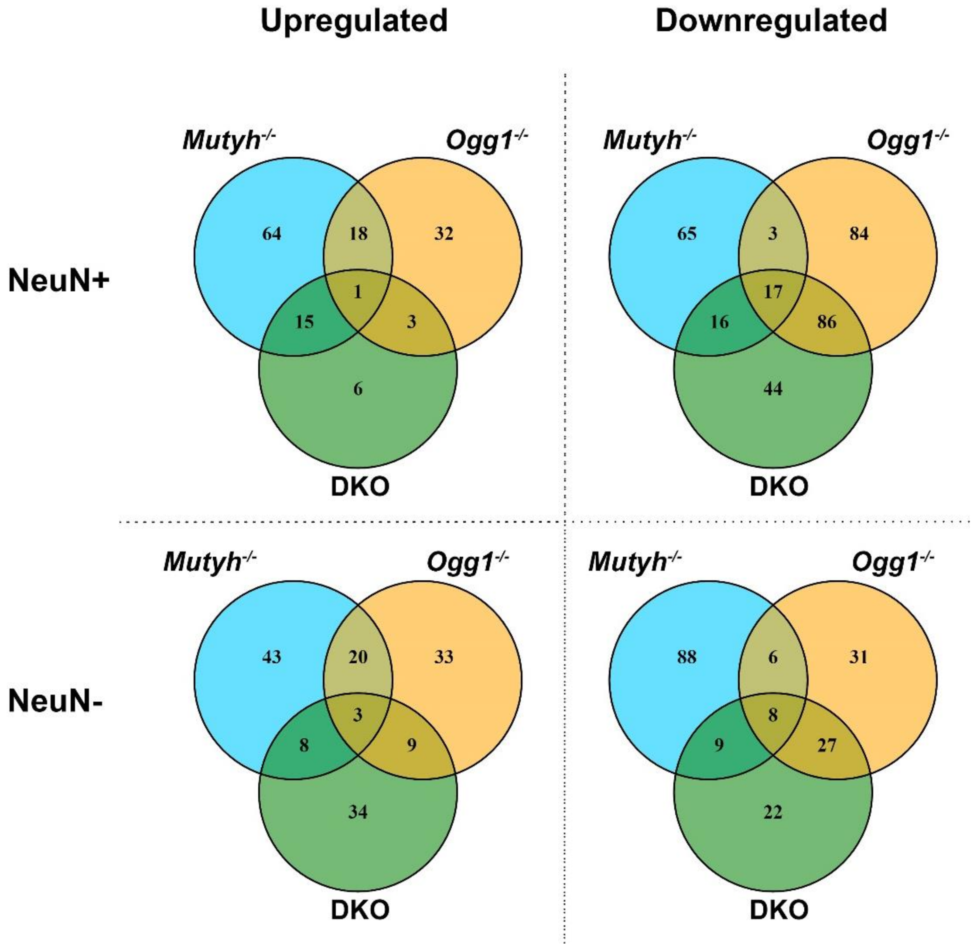


**Supplementary figure 10. Overlap of differentially expressed genes across genotypes**

Venn diagrams depict the overlap of differentially expressed genes that are either up- or downregulated among genotypes in NeuN+ or NeuN-.

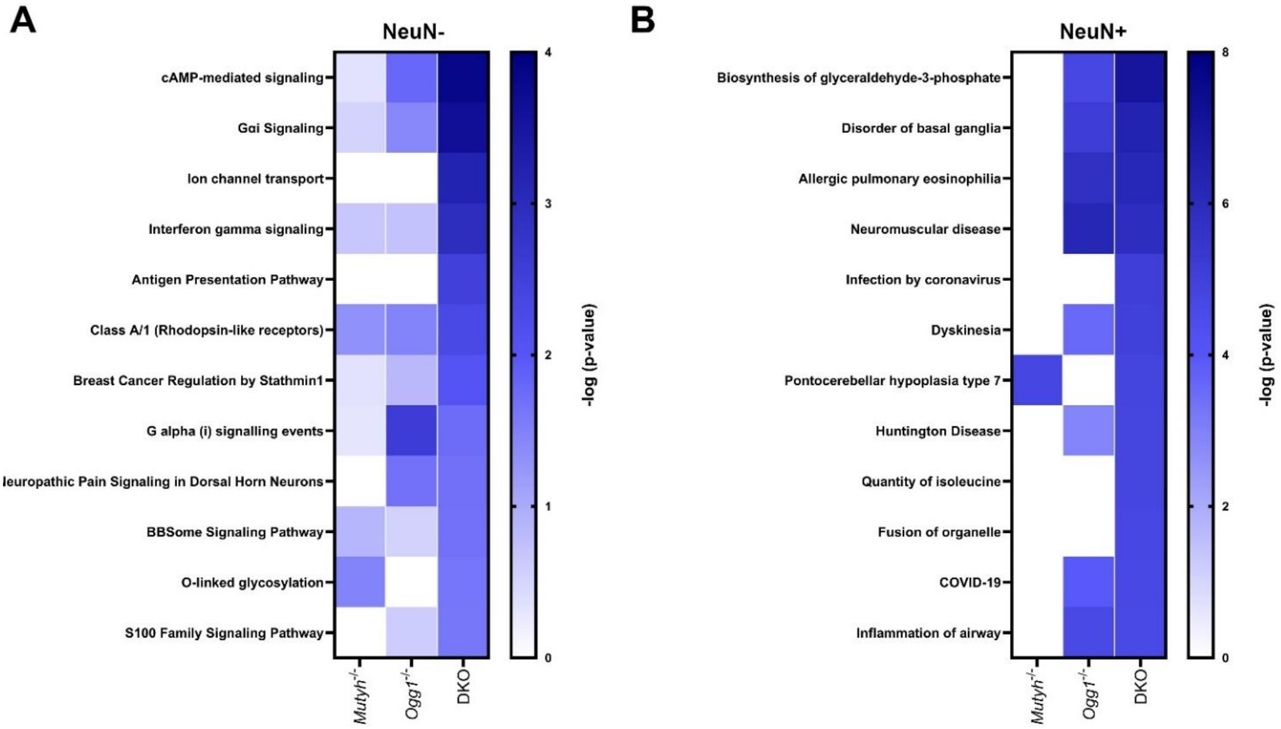


**Supplementary figure 11. Pathways analysis for differentially expressed genes**

(A) Pathways (-log10(FDR)<0.05) identified by Ingenuity pathway analysis of differentially expressed genes (DEGs) in non-neuronal cells.

(B) Disease and functions (-log10(FDR)<0.05) identified by Ingenuity pathway analysis of DEGs in neurons.

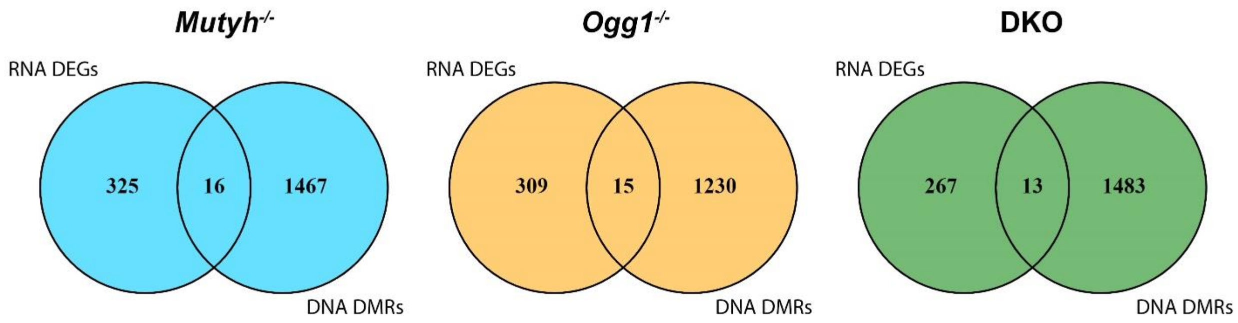


**Supplementary figure 12. Correlation between RNA expression and DNA methylation across genotypes**

Venn diagrams depict the overlap of gene expression and DNA methylation among genotypes in NeuN+ or NeuN-.
